# Self-supervised denoising for structured illumination microscopy enables long-term super-resolution live-cell imaging

**DOI:** 10.1101/2023.04.05.535684

**Authors:** Xingye Chen, Chang Qiao, Tao Jiang, Jiahao Liu, Quan Meng, Yunmin Zeng, Haoyu Chen, Hui Qiao, Dong Li, Jiamin Wu

## Abstract

Detection noise significantly degrades the quality of structured illumination microscopy (SIM) images, especially under low-light conditions. Although supervised learning based denoising methods have shown prominent advances in eliminating the noise-induced artifacts, the requirement of a large amount of high-quality training data severely limits their applications. Here we developed a pixel-realignment-based self-supervised denoising framework for SIM (PRS-SIM) that trains an SIM image denoiser with only noisy data and substantially removes the reconstruction artifacts. We demonstrated that PRS-SIM generates artifact-free images with 10-fold less fluorescence than ordinary imaging conditions while achieving comparable super-resolution capability to the ground truth (GT). Moreover, the proposed method is compatible with multiple SIM modalities such as total internal reflective fluorescence SIM (TIRF-SIM), three-dimensional SIM (3D-SIM), lattice light-sheet SIM (LLS-SIM), and non-linear SIM (NL-SIM). With PRS-SIM, we achieved long-term super-resolution live-cell imaging of various bioprocesses, revealing the clustered distribution of clathrin coated pits and detailed interaction dynamics of multiple organelles and the cytoskeleton.

## Introduction

Studying biological dynamics and functions in live cells requires imaging with high spatiotemporal resolution and low optical invasiveness. Structured illumination microscopy (SIM) is commonly recognized as a well suitable tool for live imaging because of its ability to acquire a super-resolution (SR) image from only a small number of illumination pattern-modulated images^1, 2^. However, conventional SIM reconstruction algorithm is prone to generate photon noise-induced artifacts especially under low light conditions, which substantially degrades the image quality and overwhelms useful structural information, thereby inhibiting us from fully exploring the underlying biological processs^3, 4^. To alleviate the reconstruction noise, a long camera exposure time and high excitation power are usually applied in SIM imaging experiments, which reduce the image acquisition speed and introduce considerable photobleaching and phototoxicity. This tradeoff severely limits the application of SIM in live-cell imaging.

Accompanied with the development of SIM instruments^5-7^, many techniques and algorithms aiming to reconstruct high-quality SR-SIM images with low signal-to-noise ratio (SNR) inputs have been proposed. Some algorithms have been developed to analytically improve the estimation precision of the illumination pattern^8, 9^ or iteratively denoise the reconstructed SR images under certain optical models and assumptions^10-12^. However, since the imaging process is complex and the image restoration/denoising problem is theoretically ill-posed, these algorithms cannot fully address the statistical complexity and have limited noise suppression capability^13^. Recently, deep neural networks (DNNs) have shown outstanding performance in image restoration tasks^14^. Various deep-learning-based SIM algorithms have demonstrated great potential in reconstructing high-quality SR images, even under extreme imaging conditions. Nevertheless, existing methods still face several challenges. First, some existing techniques employ “end-to-end” schemes^15-18^, which directly transform wide-filed or raw SIM images into the SR-SIM image without fully exploiting the high-frequency information modulated by the illumination pattern, i.e., the Moore fringes. As a result, the entire framework degrades to an SR inference task (termed “image super-resolution”^19, 20^) instead of analytical SR reconstruction^21^. Second, a large number of well-matched low-and high-SNR image pairs are necessary to construct the training dataset^22, 23^, which is laborious and even infeasible for biological specimens of low fluorescent efficiency or high dynamic. Third, the generalizability of the neural network is limited because in the supervised training scheme, a pre-trained denoising model cannot be reliably transferred to unseen domain with only noisy data, which inhibits the discovery of unprecedented biological structures and bioprocesses.

Here we proposed a pixel-realignment-based self-supervised method for structured illumination microscopy (PRS-SIM), which employs a deep neural network to achieve artifact-free reconstruction with ∼10 fold fewer collected photons than that used for conventional SIM algorithms^7^. The proposed PRS-SIM framework has several key advantages: first, because the analytical SIM reconstruction principle is embedded in the training and inference framework, the resolution enhancement is physically guaranteed by the SIM configuration rather than computationally achieved via data-driven supervised learning^19, 24-26^. Second, the PRS-SIM models are trained on low-SNR raw images only, without the requirement for either high-SNR ground-truth data or repeated acquisition of the same sample, resulting in a more feasible data acquisition process. Third, for time-lapse imaging, PRS-SIM can be implemented in an adaptive training mode, in which the collected low-SNR data are used to train a new customized model or fine-tune a pretrained model. Finally, PRS-SIM is compatible with multimodal SIM configurations, including total internal reflective fluorescence SIM (TIRF-SIM)^5^, grazing incidence SIM (GI-SIM)^7^, three dimensional SIM (3D-SIM)^2^, lattice light-sheet SIM (LLS-SIM)^27^, and non-linear SIM (NL-SIM)^28,29^. Benefiting from these advances, PRS-SIM instantly enables long-term volumetric SR imaging of live cells with extremely low photo-damage to the biological samples.

## Results

### The principle of PRS-SIM

The principle of PRS-SIM is schematized in Fig. 1. The PRS-SIM framework involves self-supervised neural network training (Fig. 1a) and the corresponding inference phase from raw SIM images (Fig. 1b). Specifically, the training dataset is constructed with noisy raw images based on a pixel realignment strategy, whose underlying mechanism is to utilize the similarity between adjacent pixels^30, 31^. For each noisy raw SIM image stack, we firstly applied pixel realignment strategy, which includes three operations of pixel extraction, up-sampling and sub-pixel registration (Method), to generate four raw image stacks of the same scene. Then by applying conventional SIM algorithm, four matched raw SR images are reconstructed, which are subsequently arranged as the input and target reciprocally for network training. By iteratively optimizing the L2-norm loss function, the neural network is able to transform noisy SIM images into their corresponding clean counterparts. Notably, we theoretically proved the convergence of adopting these SIM images in the loss calculation (Supplementary Note 1). In the inference phase, the raw images are firstly reconstructed into the noisy SR images via the conventional SIM algorithm, then the well-trained PRS-SIM model takes these noisy SIM images as inputs and outputs the final noise-free SR images.

**Fig. 1.**
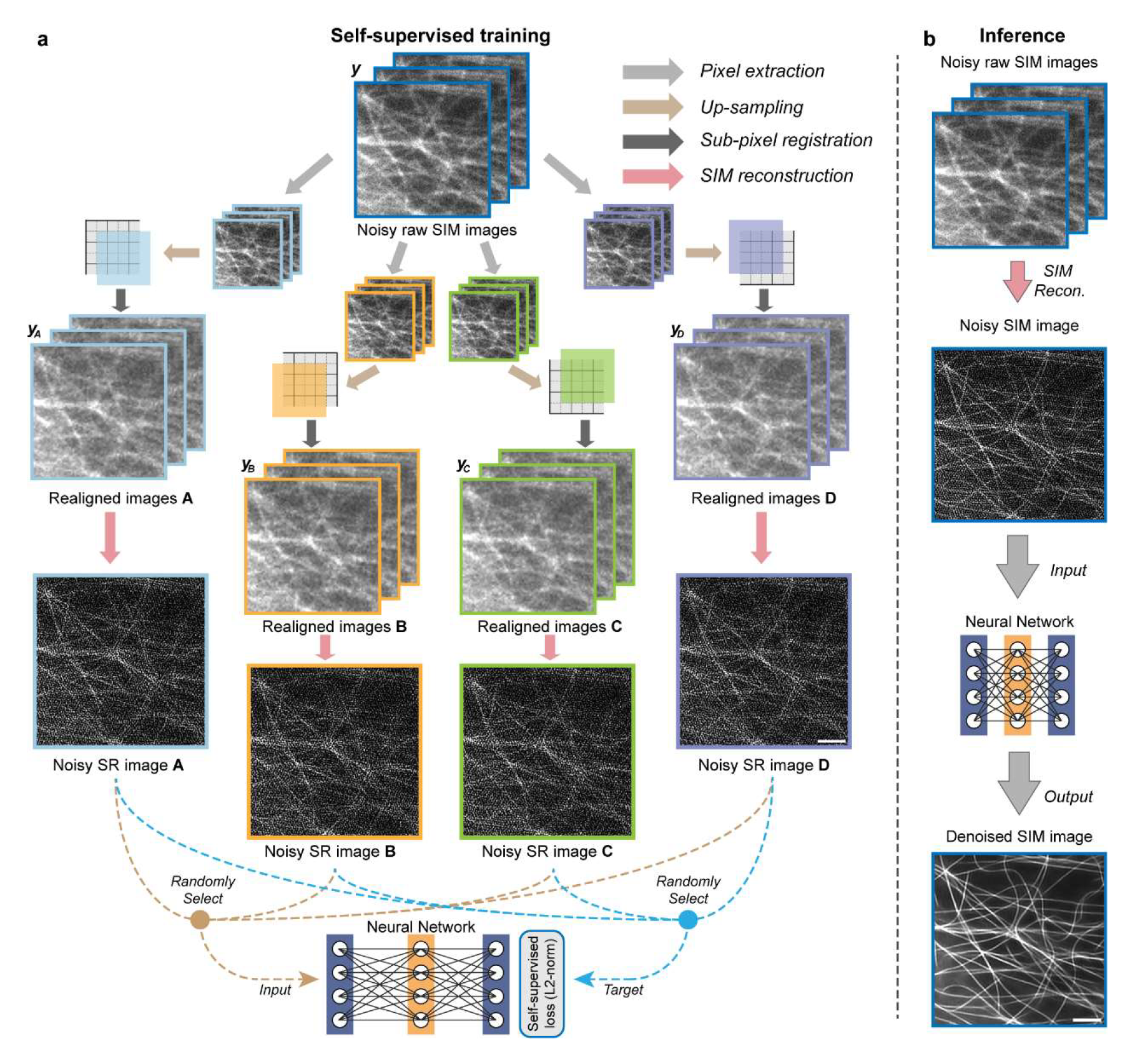
Schematic of PRS-SIM. **a**, Self-supervised training strategy of PRS-SIM. Four matched image groups ***y_A_***, ***y_B_***, ***y_C_***, and ***y_D_*** are generated by applying pixel realignment operation to a noisy low-resolution (LR) raw SIM image group ***y***. Then with conventional SIM algorithm, four super-resolution (SR) images are reconstructed, which are further randomly arranged as the input and target for neural network training. **b,** Inference pipeline of PRS-SIM. The noisy raw SIM image group are firstly reconstructed into a noisy SR image by conventional SIM algorithm. Then by inputting this noisy SR image into the pre-trained PRS-SIM model, the corresponding noise-free SR SIM image will be generated. Scale bar, 2 μm.

We first systematically evaluated PRS-SIM on the publicly available biological image dataset BioSR^16^. To quantify the performance of PRS-SIM, we calculated the peak signal-to-noise ratio (PSNR) and structural similarity (SSIM) using ground-truth (GT) SIM images as the criteria (Methods). Three individual neural networks were trained separately for clathrin-coated pits (CCPs), endoplasmic reticulum (ER), and microtubules (MTs), as representative examples of hollow, reticular, and filament structures, respectively. The training dataset was augmented with raw data from signal level 1 to signal level 4 in BioSR, and the average effective photon counts of these samples are ∼10-fold less than those used in artifact-free GT-SIM images. We compared PRS-SIM with conventional SIM (conv. SIM) and sparse-deconvolution SIM (Sparse-SIM) (Fig. 2a) and found that the detailed information can hardly be distinguished in conv. SIM and Sparse-SIM due to severe reconstruction artifacts. In contrast, PRS-SIM can clearly super-resolve ring-shaped CCPs and densely interlaced MTs, resulting in an image quality comparable to GT-SIM. The statistical results in terms of the PSNR and SSIM of 40 individual cells for each sample demonstrated that PRS-SIM achieves substantially improved denoising results for various types of specimens (Fig. 2b). The intensity profiles shown in Fig. 2c indicated that PRS-SIM successfully distinguishes several adjacent microtubules as clearly as GT-SIM, which are indistinguishable with the other methods. Furthermore, we validated the robustness of PRS-SIM on both synthetic (Supplementary Note 2) and experimental data with different signal levels and demonstrated that PRS-SIM is applicable with a wide range of input SNRs (Extended Data Figs. 1 and 2).

**Fig. 2.**
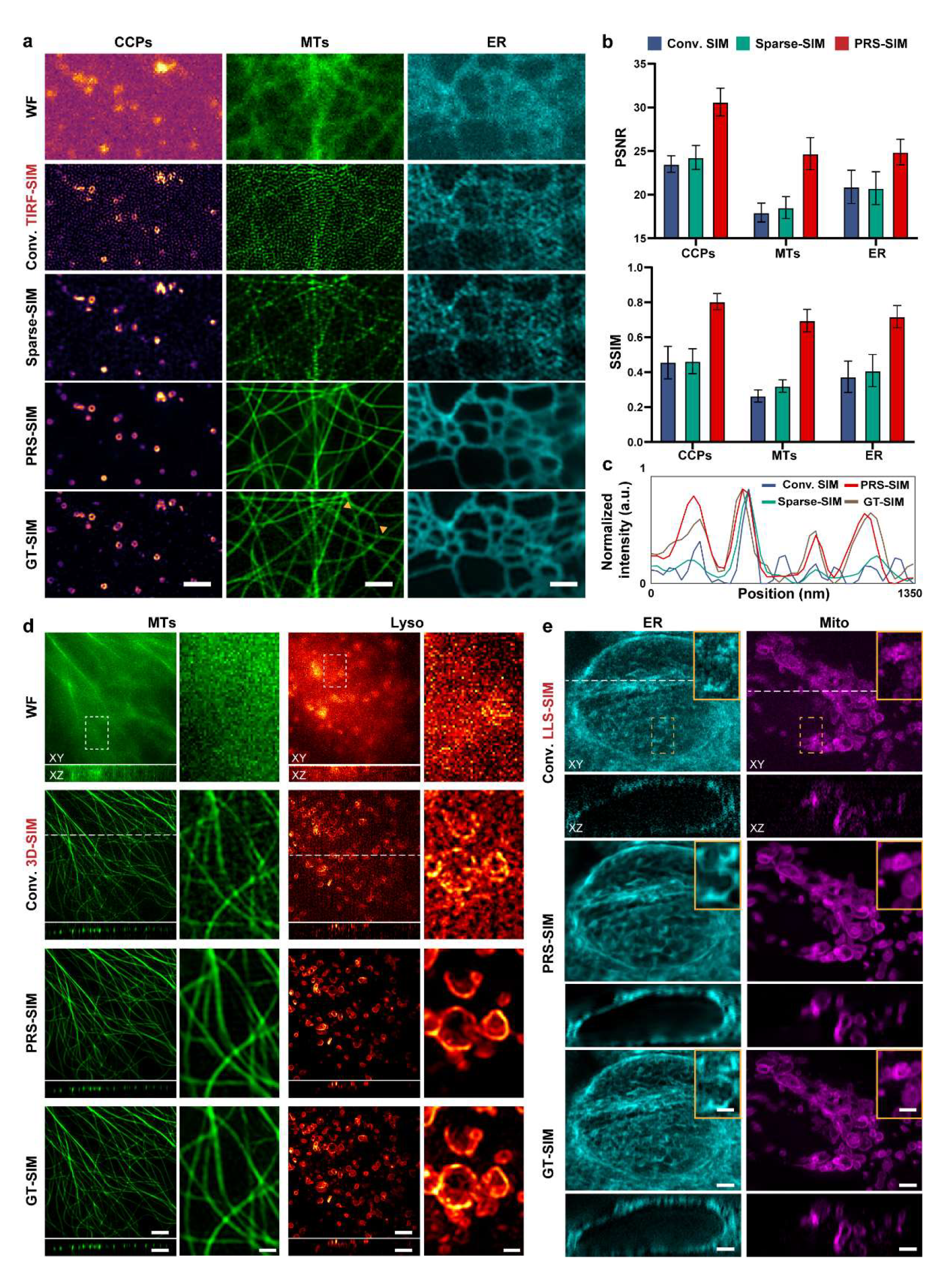
PRS-SIM on multimodal SIM systems. **a,** TIRF-SIM images of CCPs, MTs, and ER reconstructed and processed with Conv. SIM, Sparse-SIM, and PRS-SIM. WF and GT-SIM images are provided for reference. Scale bar, 2 μm. **b,** Quantitative comparison among PRS-SIM, Conv. SIM and Sparse-SIM. The PSNR and SSIM values are calculated referring to GT-SIM images (N=40 for each data point). **c,** Intensity profiles of Conv. SIM (blue), Sparse-SIM (green), PRS-SIM (red), and GT-SIM (brown) along the line indicated by the yellow arrowheads in **a**. **d,** 3D-SIM images of MTs and Lyso in fixed COS7 cells reconstructed with Conv. SIM and PRS-SIM. WF and GT-SIM images are provided for comparison. Scale bar: 1 μm, 0.5 μm (zoom-in regions) **e,** LLS-SIM images of ER and mitochondria in fixed COS7 cells reconstructed with Conv. SIM and PRS-SIM. The maximum intensity projection (MIP) of XY view and the sectioned view in XZ plane (indicated by white dashed lines in XY-views) are shown in **d** and **e**. Scale bar, 5 μm, 1 μm (zoom-in regions).

Next, we compared PRS-SIM with the classical noise2noise (N2N) method^32^, which requires two independently captured images of the same scene to train a denoiser (Methods). This requirement is impractical when the biological samples are highly dynamic or the total number of frames is limited due to photobleaching and phototoxicity. Resorting to the self-supervised training scheme, a single SIM capture for each scene is enough to train a PRS-SIM model. We compared PRS-SIM and N2N-SIM using synthetic structure with different moving speeds (Extended Data Fig. 3) and noted that as the moving speed increased, N2N-SIM generated considerably deteriorated SIM images and was prone to oversmoothing the details of subcellular structures. Compared with N2N-SIM, the proposed PRS-SIM maintained a steady denoising performance regardless of the sample moving speed, indicating the superb live-cell imaging capability especially for samples of high dynamics.

**Fig. 3.**
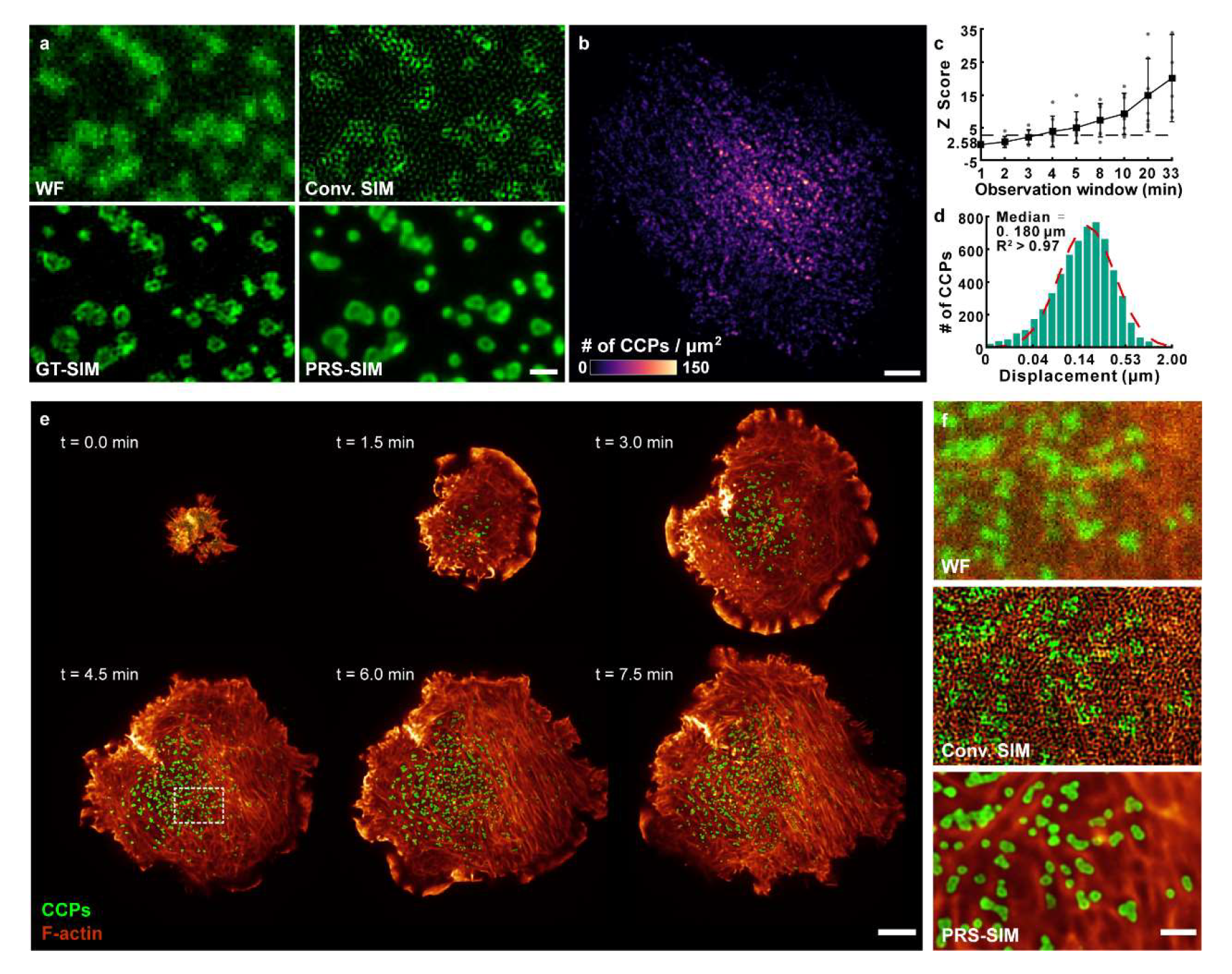
Long-term observation of the bioprocesses sensitive to phototoxicity via PRS-SIM under low excitation power. **a,** Representative PRS-SIM image (bottom right) of clathrin coated pits (CCPs) whose raw images (top left) were acquired at 30-fold lower fluorescence than those of GT-SIM (bottom left), but conveys high-fidelity ring-like structure and prevents reconstruction artifacts fulfilled in conventional TIRF-SIM image (top right). **b**, Spatial distribution of CCP nucleation events across the plasma membrane of a SUM-159 cell over 5000 frames. **c**, z-score of CCP nucleation calculated from 7 cells rapidly increases as extending the observation window. z-score gets larger than 4.95 when observation window is longer than 4 minutes, indicating that there is a less than 1% likelihood that the clustered pattern of CCPs’ nucleation could be the result of random occurrence. **d**, Histogram of mean square displacement (MSD) of 3572 CCP tracks from 3 cells. **e,** Representative frames of the dual-color time-lapse imaging of CCPs (green) and F-actin (red) in a live SUM159 cell during the growth process via PRS-SIM. The whole imaging duration is ∼8 minutes with more than 170 SR-SIM frames. **f,** Zoom-in regions (indicated by the white square in **e**) demonstrating the interaction between CCPs and F-actin. PRS-SIM (bottom) enhanced the resolution of both structures compared to WF images (top), and removed most artifacts in Conv. SIM (middle), enabling a clear visualization of the ring-like CCPs and interlaced actin filaments. Scale bar, 0.5 μm (a), 5 μm (b, e), 1 μm (f).

In addition to PRS-SIM, many other self-supervised denoising methods for fluorescence microscopy have been developed in recent years, and the blind spot-based denoising algoritm^33^ is one of the most representative approaches. Nevertheless, although these methods have shown great denoising performance for natural and microscopic images, they are not applicable to SIM images for two critical reasons. First, if the denoising algorithms are applied to raw SIM images (Extended Data Fig. 4a), i.e., images that are captured directly by the sensor, the algorithms have difficulty recognizing the illumination patterns and restoring the subtle Moiré fringes, thereby missing high-frequency information and generating reconstructed images with riddling artifacts (Extended Data Fig. 4c). Second, if these algorithms are employed in the post-reconstruction procedure (Extended Data Fig. 4b), the strongly self-correlated noise patterns in the reconstructed SIM images are inconsistent with the blind-spot principle, leading to poor denoising performance. The proposed PRS-SIM scheme addresses these issues by leveraging the intrinsic linearity of SIM reconstruction and integrating this physical property into the objective function design, thereby yielding superior restoration capability for SIM images. We experimentally compared PRS-SIM models with two representative self-supervised denoising approaches: noise2void (N2V)^33^ and hierarchical diverse denoising (HDN)^34^. Both the perceptual comparisons and the quantitative analysis showed that PRS-SIM can generate SR images with considerably fewer artifacts, outperforming other self-supervised denoising methods by a large margin (Extended Data Fig. 5).

**Fig. 4.**
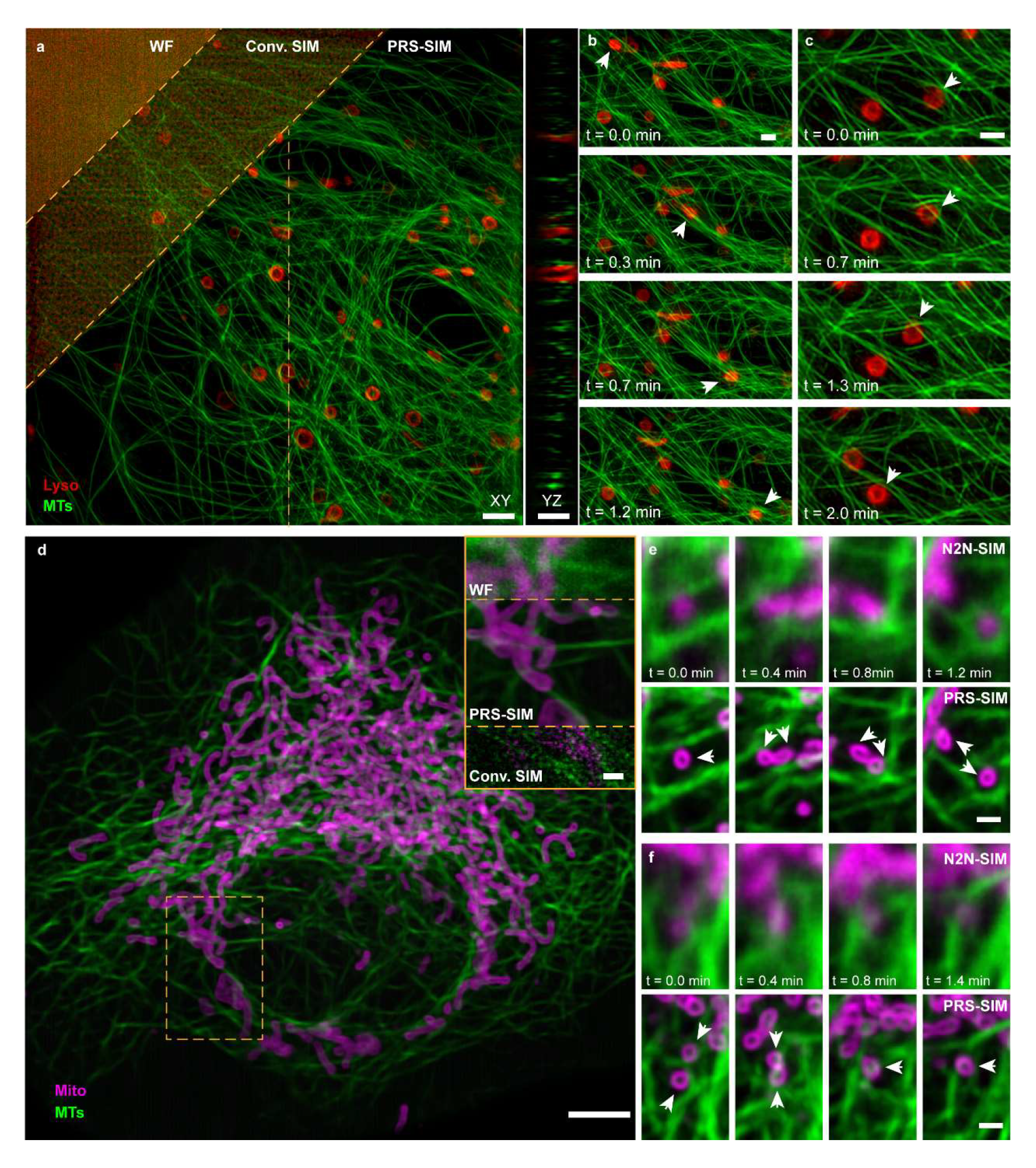
Long-term volumetric super-resolution imaging of live cells with adaptively trained PRS-SIM. **a,** Progression of resolution and quality improvement of a live COS7 cell expressing 3xmEmerald-Ensconsin (green) and Lamp1-Halo (red), from wide-field, Conv. 3D-SIM, and PRS-SIM enhanced 3D-SIM. **b, c**, Time-lapse PRS-SIM images of lysosomes moving along adjacent MTs (**b**) or deforming under the traction of MTs (**c**) as indicated by white arrows. **d**, Representative PRS-SIM enhanced LLS-SIM images of a live COS7 cell expressing TOMM20-2xmEmerald (magenta) and 3xmCherry-Ensconsin (green). The comparison of WF, Conv. SIM and PRS-SIM images of a zoom-in region are displayed in the top right corner. **e, f**, Time-lapse recordings of the fission (**e**) and fusion (**f**) processes of mitochondria under the interaction with MTs as indicated by white arrows. The denoising results of N2N-SIM and PRS-SIM are compared to demonstrate their performance on fast-moving samples. Both the adaptively trained PRS-SIM models and N2N-SIM models were trained only with the noisy raw time-lapse data. Scale bar, 5 μm (a, d), 1 μm (b, c, e-f, and zoom-in regions of d).

Due to the internal similarity of the post-processing pipeline for various SIM modalities, besides TIRF/GI-SIM, PRS-SIM is compatible with other SIM configurations such as NL-SIM (Extended Data Fig. 6), 3D-SIM, and LLS-SIM for higher resolution or volumetric SR imaging under low-light conditions. For 3D-SIM, we evaluated the performance of PRS-SIM by processing the images of microtubules labelled with 3xmEmerald-Ensconsin and lysosomes (Lyso) labelled with Lamp1− mEmerald in fixed COS7 cells (Fig. 2d and Extended Data Fig. 7). For each sample, ∼20 individual cells were imaged under low and high illumination conditions to acquire noisy data and the corresponding high SNR reference, respectively. The raw SIM data were first reconstructed into 3D SR volumes via the conventional 3D-SIM algorithm and then denoised with 3D PRS-SIM models, which were modified into 3D U-net^35^ architectures from the original 2D version (Methods) and trained with the noisy data only.

The orthogonal view of the representative PRS-SIM images indicated that most of the noise-induced artifacts in the conventional 3D-SIM results were removed by PRS-SIM, and the reconstruction quality of PRS-SIM is comparable to that of GT-SIM in both the XY plane and Z-axis (Fig. 2d). For the LLS-SIM configuration, we employed our home-built LLS-SIM system to acquire raw images of mitochondria (Mito) labelled with TOMM20-2xmEmerald and endoplasmic reticulum labelled with calnexin-mEmerald following a similar procedure as 3D-SIM. For both structures, PRS-SIM achieved a substantial improvement in both perceptual quality and statistical metrics (Fig. 2e and Extended Data Fig. 8) across a field-of-view (FOV) of 70μm × 47μm × 27μm (after de-skewing). These results suggest that PRS-SIM shows a great potential for extending the application scope of multimodal SIM to low-light conditions without the need to acquire abundant training data.

### Observation of bioprocesses sensitive to phototoxicity

One major limitation of SIM is the requirement of high-intensity illumination, resulting in substantial phototoxic side effects. This phototoxicity largely limits the SR imaging duration for live specimens, particularly when imaging molecules with low expression levels or processes that are vulnerable to high-dose illumination. To demonstrate the potential of our method in reducing the required light dose, we first applied PRS-SIM to visualize clathrin-mediated endocytosis in gene-edited SUM159 cells expressing clathrin-EGFP at endogenous levels. The limited fluorescence of these cells prevents conventional TIRF-SIM (conv. TIRF-SIM) imaging from more than 150 frames, corresponding to an imaging time of ∼3 minutes^6^, because under low SNR conditions, conv. TIRF-SIM image contained substantial reconstruction artifacts (Fig. 3a). Although the fluorescence intensity of each raw image was 30-fold less than that of the high-SNR GT-SIM image, PRS-SIM was still able to reconstruct high-fidelity SR information of the hollow, ring-like structure of CCPs (Fig. 3a). Therefore, PRS-SIM allowed us to characterize clathrin-mediated endocytosis at high spatiotemporal resolution for an unprecedented imaging duration of more than 5,000 frames, corresponding to an imaging time of more than 45 minutes. Previous studies have reported that clathrin-mediated endocytosis is initiated randomly based on analyses of the distribution of all CCP nucleation events over the limited observation window of ∼7 minutes^36, 37^. By imaging the same process over 45 minutes, we found that most CCP nucleation sites tended to be spatially clustered (Fig. 3b, c, z-score > 20, n = 7 cells; Methods), with many events occurring in confined regions, possibly at stable clathrin coated plaques^38^. Moreover, after tracking the CCP trajectories from their initiation to their detachment from the plasma membrane, we noted that the displacement of most CCPs was relatively small (Fig. 3d, Median = 0.180 μm). This finding is consistent with clathrin uncoating occurring near the site of invagination of the coated pit.

We also utilized PRS-SIM to investigate dynamic interactions between subcellular organelles and the cytoskeleton in SUM159 cells. Since the growing cells are light-sensitive and fragile, we decreased the illumination power to 10% of that used for usual experiments to image the entire adhesion process after dropping a SUM159 cell onto a coverslip. Under the low excitation intensity conditions, we successfully recorded the detailed interactions between CCPs and F-actin during the cell adhesion and migration for ∼8 minutes with more than 170 SR-SIM frames (Fig. 3e). As shown in Fig. 3f, the hollow structure of CCPs (green) and the densely interlaced F-actin (orange) cannot be resolved in wide-field (WF) and conventional SIM (conv. SIM) images due to the diffraction limitation in WF microscopy and noise-induced artifacts in conv. SIM images. In contrast, the fine structures of CCPs and F-actin were both clearly distinguished by PRS-SIM, enabling further study of their detailed interactions. We next applied the Weka segmentation algorithm to extract the filament skeleton and calculated the Mander’s overlap coefficient (MOC) between the two structures in each frame (Methods; Extended Data Fig. 9). We found that the MOC remained in a relatively small value during the whole adhesion process, indicating that most CCPs stayed at the interspace of actin filaments and were intensively regulated by the cytoskeleton throughout the adhesion process.

### Long-term volumetric SR imaging of subcellular dynamics with adaptive trained PRS-SIM

Volumetric SIM imaging, such as 3D-SIM and LLS-SIM, causes severer photo-damage to live specimens than 2D-SIM (TIRF-SIM)^12^. To realize long-term volumetric SR live-cell imaging, we equipped our multi-SIM system with PRS-SIM and imaged a live COS7 cell expressing 3xmEmerald-Ensconsin (green) and Lamp1-Halo (red) in 3D-SIM mode under ∼10-fold lower excitation power than typical imaging conditions (Fig. 4a-c). The data were acquired over 1 hour (400 two-color SIM volumes at an interval of 10 seconds). During the data acquisition process, no decrease in cell activity was observed, indicating negligible phototoxicity effects. Although conventional SIM reconstruction reduces the out-of-focus fluorescence and improves the axial resolution, the detection noise severely degrades the image quality, preventing us from investigating the underlying bioprocesses. In contrast, the PRS-SIM model, which was trained by only 30 selected frames with the equal distance from the noisy time-lapse data, substantially removed the reconstruction artifacts and restored the fine structures of both organelles including continuous microtubule filaments and the hollow lysosomes. These advantages of PRS-SIM enable a clear volumetric observation of the dynamic interaction between microtubules and lysosomes, e.g., the directional movement of a lysosome along the MT filaments (Fig. 4b) and the hitchhiking remodeling mechanism of MT filaments under the traction of lysosomes (Fig. 4c).

We next applied the PRS-SIM enhanced LLS-SIM system to record the volumetric subcellular dynamics of COS7 cells expressing TOMM20-2xmEmerald and 3xmCherry-Ensconsin (Fig. 4d-f). Two PRS-SIM models for mitochondria (Mito) and MTs were independently trained with the noisy time-lapse data themselves, which consisted of ∼310 two-color SIM volumes acquired at an interval of 12 seconds. We demonstrated that the adaptively trained PRS-SIM models removed most noise-induced artifacts and resolved the delicate structures of Mito and MTs (Fig. 4d). However, due to the rapid movement and deformation of the two observed structures, the classical denoising algorithm N2N^32^ and its derivative DeepCAD^39, 40^, which are based on the temporal continuity between adjacent frames (Methods), generated oversmoothed images with severe motion blur (Fig. 4e-f, Extended Data Fig. 10). With the prolonged observation window provided by PRS-SIM, we clearly identified the fission and fusion processes of Mito (Fig. 4e, f), which are some of the most common yet very important bioprocesses in live cells. Moreover, we emphasized that since the adaptive training mode of PRS-SIM utilizes only the noisy collected data for network training and then denoises themselves, there is no domain shift problem. Thus, the adaptively trained PRS-SIM models provide a high denoising fidelity and show great potential in the discovery of previously unseen biological structures and phenomena.

## Discussion

In summary, PRS-SIM is a novel self-supervised learning-based method for SIM image restoration, which trains the denoiser with only noisy data and reconstructs artifact-free SR-SIM images with 10-fold less fluorescence than routine SIM imaging conditions. The proposed self-supervised strategy does not require either high-SNR GT data or repeated acquisition to construct the training dataset. Thus, this easy-to-implement data acquisition scheme is applicable to biological specimens of high dynamics or with low fluorescence efficiency. For long-term live-cell imaging, PRS-SIM can be applied in the adaptive training mode, where the acquired noisy data are directly used to train the denoising model. Therefore, no pre-trained models for the same samples are needed, and with this advance, PRS-SIM can be used to discover previously unknown biological structures and phenomena. Finally, we emphasize that our method is applicable to multiple SIM modalities, including TIRF/GI-SIM, 3D-SIM, LLS-SIM, and even NL-SIM. With PRS-SIM, we achieved long-term live observations of subcellular dynamics and diverse bioprocesses with extremely low invasiveness, demonstrating the broad applicability of our method. Furthermore, to make PRS-SIM more accessible for biological research, we developed an easy-to-use Fiji toolbox^41^ (Supplementary Note 3, Supplementary Fig. 3-4), where the network training and inference can be implemented by several clicks.

PRS-SIM can be improved in several ways. First, successful PRS-SIM reconstruction relies on accurate estimations of the SIM patterns, which is challenging under extremely low-light conditions for conventional SIM parameter estimation algorithm. Therefore, an additional neural network for more precise parameter estimation may improve the robustness of PRS-SIM. Second, to obtain volumetric images of thick samples, although the noise-induced artifacts are mitigated by PRS-SIM, the image quality suffers from sample-induced optical aberrations. Incorporating PRS-SIM into an adaptive optics-embedded SIM system^42, 43^ may greatly improve the fidelity of the reconstructed SR images.

## Methods

### Optical setup

All the experiments in this work were performed on our home-built multi-modality SIM system (Multi-SIM) or lattice light-sheet SIM (LLS-SIM) system, which were developed based on previous setups^6, 7^. Three modes of TIRF-SIM, GI-SIM, and 3D-SIM were embedded in the Multi-SIM system. Briefly, three laser beams of 488 nm (Genesis-MX-SLM, Coherent), 560 nm (2RU-VFL-P-500-560, MPB Communications), and 640 nm (LBX-640-500, Oxxius) were collimated for multi-channel excitation and controlled by an AOTF (AOTFnC-400.650, AA Quanta Tech) for rapid switching. The structured illumination patterns were generated by a ferroelectric spatial light modulator (SLM, QXGA-3DM, Forth Dimension Display) placed conjugated to the sample plane. Illumination patterns of 3-phase×3-orientation for TIRF/GI-SIM mode and 3-phase×5-orientation for 3D-SIM mode were generated in our experiments. The final images were collected by an sCMOS camera (Hamamatsu, Orca Flash 4.0 v3).

For the LLS-SIM system, three laser beams of 488 nm, 560 nm and 640 nm (MPB Communications) were used for multi-color excitation. The illumination pattern is displayed on the SLM (the same type as used in Multi-SIM) and then filtered by an annular mask of an outer NA of 0.5 and inner NA of 0.375 to obtain a balanced axial and lateral resolution. A pair of galvo mirrors (Cambridge Technology, 6210H) was set for x-axis and z-axis scanning. The emission fluorescence was collected by a water-immersion objective (Nikon, CFI Apo LWD 25XW, 1.1NA) and captured by a sCMOS camera (Hamamatsu, Orca Fusion). The illumination patterns of 3-phase×1-orientation were generated for each z-slice. The oblique angle between the illumination path and the detection path is 30°. All the equipment was synchronized by a DAQ card, allowing the maximum imaging speed at ∼1000 z-slices per second. The pixel size of the detected image is 92.6 nm and the axial step size is determined by the specific scanning angle step used for each experiment.

### Data acquisition

The experiments in this work can be categorized as fixed sample imaging and time-lapse live-cell imaging. For fixed sample imaging, we utilized the data from the open-source dataset BioSR^16^ or acquired via our home-built SIM systems. For TIRF-SIM experiments, The CCPs, ER, and MTs images whose signal levels range from 1 to 4 in BioSR were used to create the training dataset. For 3D-SIM and LLS-SIM experiments, the dataset used for both training and inference was acquired with our home-built Multi-SIM and LLS-SIM systems. Specifically, for each type of specimen, we acquired more than 20 sets of raw SIM images at four escalating levels of excitation light intensity to create the training dataset, and then tuned the laser power to the maximum to capture the high-SNR images as the corresponding GT data. Notably, the training dataset is generated purely with the low-SNR data, and the high-SNR GT data are only used as the reference for quantitative analysis.

For time-lapse imaging, the 2D and 3D experiments were carried out with the TIRF-SIM and 3D-SIM mode of the Multi-SIM system, respectively. The excitation light power used in all live experiments was set to 10-fold lower than that used in common imaging conditions, corresponding to an average photon count of 40∼60 for each raw SIM image, to minimize the phototoxicity and photobleaching effects. The specific imaging conditions for each time-lapse experiment were listed in Supplementary Table 1.

### Pixel realignment strategy

The self-supervised training dataset was generated with the pixel realignment strategy. The raw dataset consists of a series of low-SNR raw SIM image groups. Each individual image in a group is a WF image under a specific illumination pattern (e.g. 3-orientation × 3-phase for 2D/TIRF-SIM and 3-orientation × 5-phase × Z-slice for 3D-SIM). For each raw SIM image group, the generation of the training dataset of PRS-SIM models mainly takes the following steps:

i. Each raw image is divided into 4 sub-images by applying a 2 × 2 down-sampler and formed four sub-image groups.
ii. The augmented four sub-image groups are re-up sampled into the original size with the nearest interpolation.
iii. Based on the position of the valid pixel in each 2 × 2 cell, a sub-pixel translation is applied to each raw image, which guarantees that they are well spatially calibrated with each other.
iv. The generated sub-images groups are reconstructed into four noisy SIM images by applying the conventional SIM algorithm.
v. Then several image patched pairs are augmented by randomly selecting two out of four noisy SIM images as the input and target, and applying ordinary data augmentation operations, e.g., random cropping, flipping and rotation.

Note that for 3D-SIM stacks, both the down-sampling, up-sampling and translation operations in step (i)-(iii) are implemented in a slice-by-slice manner. By applying the pixel realignment strategy to all noisy SIM image groups, the complete training dataset is generated. Typically, no fewer than 10 individual image groups are adequate for training a robust PRS-SIM model (Supplementary Fig. 1).

### Network architecture

PRS-SIM employs U-net^35^ as the backbone architecture, which has already shown superior performance in denoising task elsewhere^32^ (Supplementary Fig. 2). The network is composed of an encoder module and a decoder module. For the encoder module, the input data is firstly fed into a convolutional layer with 48 kernels and then encoded by five consecutive encoding blocks. Each encoding block consists of a convolutional layer followed by a non-linear activation layer and a max-pooling layer for spatial down sampling. For the decoder module, five decoding blocks are involved, each of which consists of two consecutive convolutional layer and a nearest interpolation layer for spatial up sampling. Skip-connections were embedded between the encoding and decoding blocks to prevent over-fitting. Two additional convolutional layers were placed at the end of the network to transfer the final denoised image into the same shape as the input image. Concretely, the kernel size of all the convolutional layers is 3 × 3 and the activation function used is Leaky-ReLU, which is defined as:

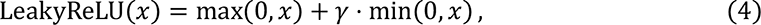

where *γ* denotes the negative slope coefficient (set as 0.1 in our experiments). For 3D-SIM applications, all the convolutional layers and pooling layers were replaced with the corresponding 3D versions and the other parts remained unchanged.

### Data processing and Network training

The training dataset of PRS-SIM consist of a series of image pairs generated only from the low-SNR raw images as described in the previous section. For pre-trained PRS-SIM models, 20-40 distinct ROIs of each type of specimens were imaged to create the training dataset. For adaptive training mode of PRS-SIM, ∼100 frames/volumes were randomly selected from the entire time series for training. Image augmentation operations, including random cropping, rotation, and flopping, were further employed to create ∼100000 mini-patch pairs of 128×128 pixels (64×64×8 voxels for 3D-SIM) to avoid overfitting.

During the network training, Adam optimizer with an initial learning rate of 10^-4^ was adopted to accelerate the convergence. A multi-step scheduler was employed to decrease the learning rate by a factor of 0.5 at the designated epochs. The training processes were performed on a workstation equipped with a graphics processing unit (Nvidia GeForce RTX 3090Ti, 24GB memory). The source codes were written based on PyTorch v1.5 framework in Python v3.6. The typical training time for a dataset of ∼100000 mini-patch pairs is about 2 hours for 2D batches and 4 hours for 3D batches. More training details of each experiment performed in this work were listed in Supplementary Table 2.

For the inference phase, the noisy raw SIM images were reconstructed into SR images via conventional SIM algorithm, divided into several tiled patches of 256×256 pixels with 10% overlap, fed into the pre-trained network, and finally stitched together to form the denoised SR images. For adaptive training mode of PRS-SIM, the time-lapse data was denoised with the model trained by itself, while in other experiments the data was denoised with the pre-trained network of the same type of specimens.

For N2N-SIM training in Fig. 4e-f, Extended Data Fig.3, and Extended Data Fig.9, we randomly selected two consecutive frames/volumes in the time-lapse data used as the input and target, respectively. The whole training dataset are generated from ∼100 independent frame/volume pairs. Other operations and configurations during training and inference are the same as PRS-SIM.

### Image assessment metrics

To quantitatively evaluate the denoised images output by PRS-SIM, we employed the peak signal-to-noise ratio (PSNR) and structural similarity (SSIM) between the denoised image *I* referring to the GT image *I_gt_* as the metric. Since the signal intensity of the denoised and GT images is of different dynamic range, we first applied percentile normalization to *I* and *I_gt_* as:

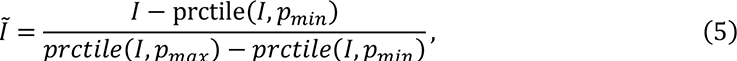

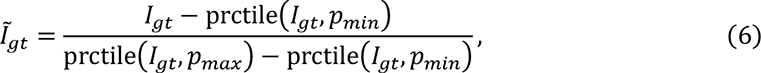

where *prctile*(*I*, *p*) denotes the intensity of the pixel ranking at *p*% of image *I*, and 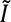 denotes the corresponding normalized image. The *p_min_* and *p_max_* are set as 0.1 and 99.5 in our analysis. To further alleviate the disturbance in metric calculation, we implemented a linear transformation to the normalized image 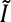 by:

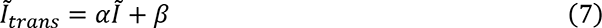

where *α* and *β* denote the transformation coefficients to minimize the square root error between the transformed image and the normalized GT image, which can be formulated as a linear regression problem:

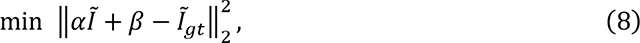

where ‖·‖_2_ is the L2-norm. The closed solution of this problem is:

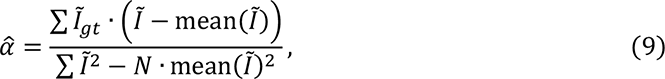

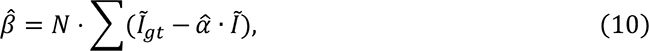

where *N* is the pixel number of the image, ∑· denotes the pixel-wise sum, â and 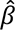 denote the optimal values of the transformation coefficients *α* and *β*, respectively. Then the final PSNR and SSIM are calculated as:

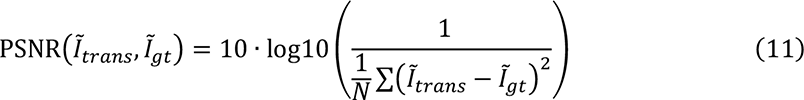

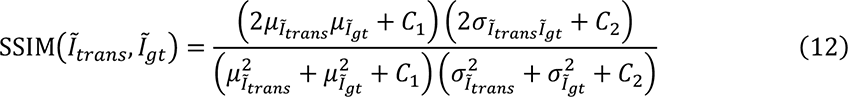

where 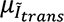, 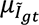 and 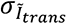, 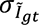 denote the mean values and standard deviations of image 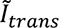 and 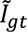, respectively; 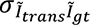 denotes the cross-covariance between 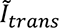 and 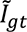. The constant *C*_1_ and *C*_2_ used in this paper is 0.01^2^ and 0.03^2^, respectively.

To characterize the resolution of the images output by PRS-SIM, we employed single-image based Fourier ring correlation (FRC) method^44^. The raw image *I* is split into two sub-images *I*_1_ and *I*_2_ by interleaved pixel extraction. Then the FRC value of the central ring region with radius *R* is calculated as:

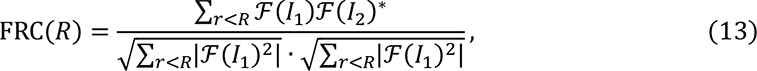

where the symbol 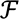 denotes Fourier transformation. By calculating the FRC value from 0 to *R_max_* (the reciprocal of the pixel size), a generally declining curve is formulated. The resolution can be measured as the reciprocal of the Fourier cutoff frequency *R_cutoff_*, where FRC(*R_cutoff_*) < *tsh*, where *tsh* represents the spectral intensity threshold. In our analysis, the *tsh* is set as a typical value of 0.25.

### Data analysis

We utilized the spatial autocorrelation (*i.e.*, Global Moran’s Index^45^) to evaluate whether the distribution of clathrin coated pit (CCP) nucleation sites is clustered, dispersed, or random. For each time-lapse dataset, we first localized the centroid positions of all CCPs at each time point, and then linked them temporally in the whole time series using the ImageJ plugin TrackMate^46^, thus yielded trajectories of all detected CCPs. To rule out the false-positive events, the trajectories of less than 40 time points corresponding to a duration of 20 seconds were excluded from following computation. Subsequently, for each time-lapse data, the initial locations of the CCP trajectories detected in the designated observation window were projected onto the same image as the CCP nucleation sites’ map (Fig. 3b). Then, the Moran’s Index can be calculated as:

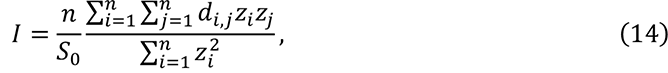

where 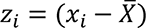 is the deviation of the event count of the *i^th^* pixel from the average count; *d_i,j_* refers to the inverse Euclidean distance between pixel *i* and *j*; *n* is the total pixel number of the map and 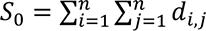 is the summation of *d_i,j_*. Finally, the *z*-score was calculated for each nucleation sites map to evaluate the significance of the Moran’s Index (Fig. 3c):

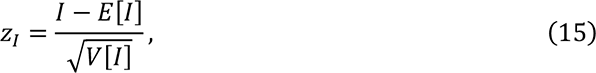

where *E*[·] and *V*[·] are the expectation and the variance of *I*, respectively. In general, the larger *z*-score indicates the stronger tendency of clustering.

To quantitatively investigated the interaction of organelles during the cell growth (Fig. 3e, f, Extended Data Fig. 8), we calculated the Mander’s overlapped coefficient (MOC)^47^ of CCPs referring to F-actin. For each frame, a binary mask (denoted as *M*) is firstly generated by applying a threshold *tsh_M_* to the F-actin channel, which represents the F-actin skeleton:

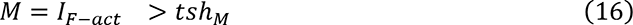

Then the *MOC* value is calculated as:

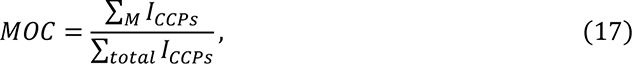

Denote ∑*_M_I_CCPS_* the ∑_*totla*_ *I_CCPS_* intensity summation of the CCP channel within the masked region and the entire image, respectively.

### Cell culture, transfection, and stain

Cos7 cells were cultured in DMEM (Gibco), supplemented with 10% fetal bovine serum (Gibco) and 1% penicillin-streptomycin in 37℃ with 5% CO_2_. For live cell imaging, the coverslips were pre-coated with 50μg ml^-1^ of collagen and cells were seeded onto coverslips with about 70% density before transfection. After 12h, cells were transfected with plasmids using Lipofectamine 3000 (Invitrogen) according to the manufacturer’s protocol. Cells were imaged 12-24 hours after transfection in a stage top incubator (Okolab) to maintain condition at 37℃ with 5% CO_2_. The plasmid constructs used in this work were 3xmEmerald-Ensconsin, Lamp1-mEmerald, TOMM20- 2xmEmerald, calnexin-mEmrald, and Lamp1-Halo.

SUM159 cells were genome edited to incorporate EGFP to the C-terminus of clathrin light chain A (clathrin-EGFP) using the TALEN-based approach^48^. The clathrin-EGFP expressing cells were enriched by two sequential bulk sorting. The cells were cultured in DMEM/F-12 (Gibco) medium supplemented with 5% fetal bovine serum (Gibco), 5 μg/ml Bovine insulin (Cell Applications), 10 mM HEPES (Gibco), 1 μg/mL Hydrocortisone (Sigma) and 1% Penicillin-Streptomycin (Gibco) in 37℃ with 5% CO2. For dual-color experiments, these SUM-ki-CLAT-GFP cells were further transfected with the lifeact-Halo. Before imaging, we digested the cells using 0.25% Trypsin, and then dropped cell suspension onto the coverslip pre-treated with 50 μg/mL collagen.

## Acknowledgement

This work was supported by the National Natural Science Foundation of China (31827802, 32125024, 31970659, 32271513, 62071271, 62088102, and 62222508); the Ministry of Science and Technology (2021YFA1300303, 2020AA0105500); China Nation Postdoctoral Program for Innovative Talents (BX2021159); Shuimu Tsinghua Scholar Program (2021SM039, 2022SM035), China Postdoctoral Science Foundation (2022M721842); Beijing Natural Science Foundation (JQ21012); the Tencent Foundation through the XPLORER PRIZE; and the Youth Innovation Promotion Association of the Chinese Academy of Sciences (2020094).

## Extended Data Figures

**Extended Data Fig. 1.**
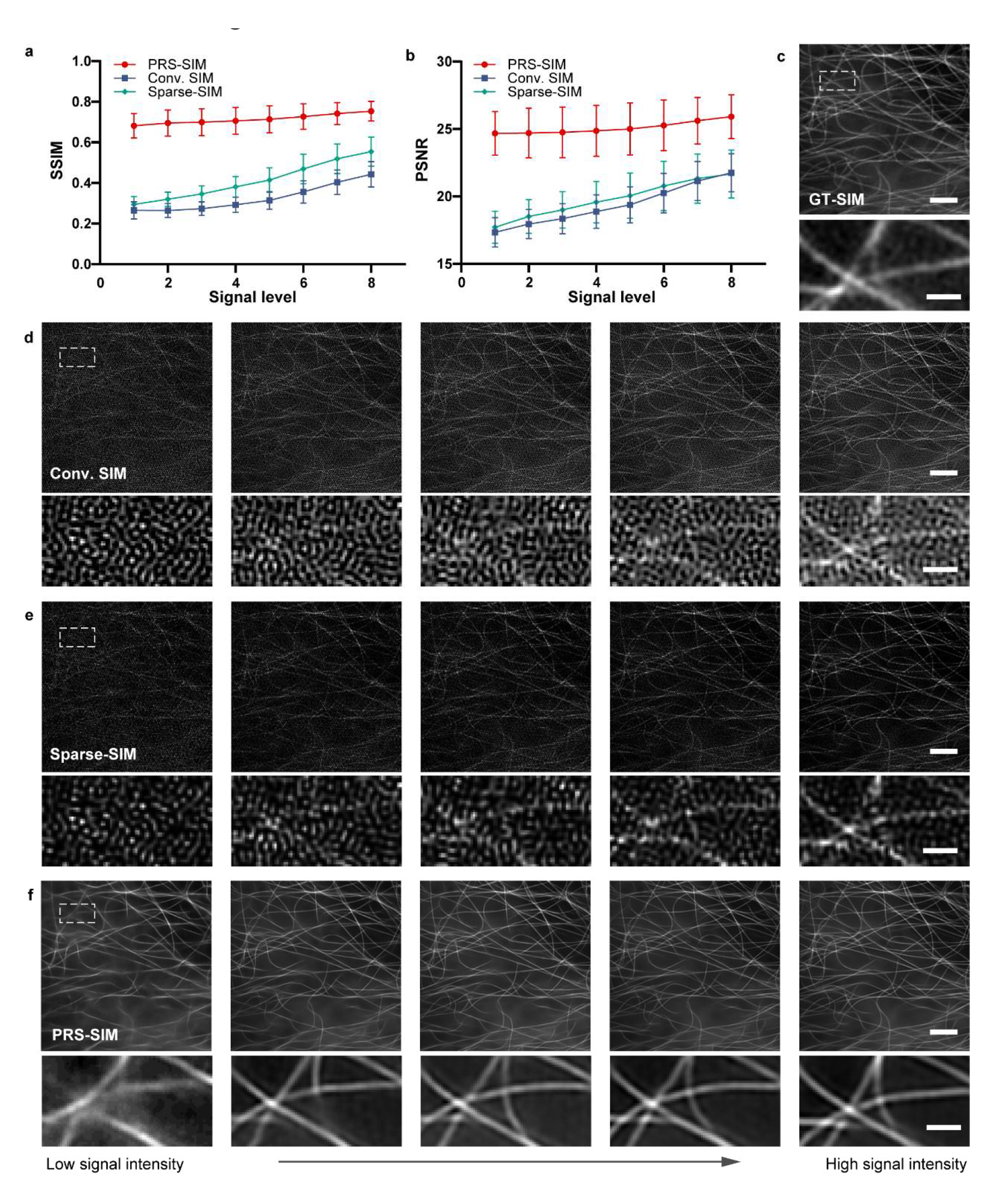
Evaluation of PRS-SIM on experimental data with different input signal levels. **a,b,** The SSIM (**a**) and PSNR (**b**) evaluation of PRS-SIM, Conv. SIM and Sparse-SIM referring to GT-SIM over different input signal levels (level 1-8 from the publicly accessible dataset BioSR). Sample size: N=40 for each data point. **c,** Representative GT-SIM image of MTs. **d-f,** SR images reconstructed via Conv. SIM (**d**), Sparse-SIM (**e**), and PRS-SIM (**f**) from different input signal intensities. Both the quantitative analysis and the reconstructed images demonstrated that PRS-SIM has substantially better performance than Conv. SIM and Sparse-SIM, and is capable of removing the noise-induced artifacts over a wide range of input signal intensities. Scale bar, 2 μm (regular), 0.5 μm (zoom-in).

**Extended Data Fig. 2.**
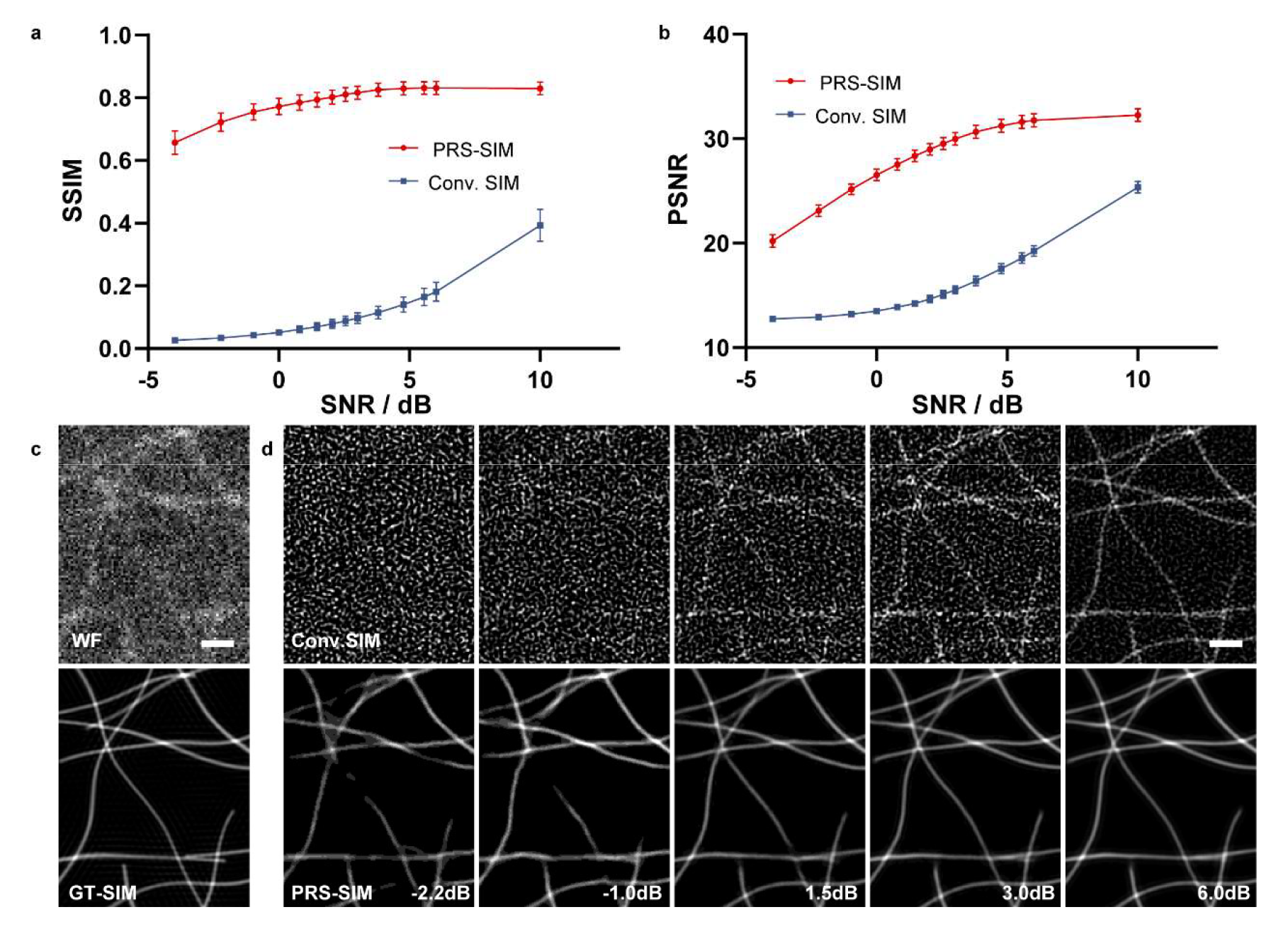
Performance validation of PRS-SIM on synthetic filaments. **a,b,** The SSIM (**a**) and PSNR (**b**) comparisons of PRS-SIM and Conv. SIM referring to GT-SIM over different SNRs of raw data. Sample size: N=50 for each data point. **c,** The representative noisy WF images and GT-SIM images. **d,** The Conv. SIM (upper) and PRS-SIM (lower) images of the ground-truth (**c**) under different SNRs. Both the quantitative and visualization results demonstrated the significant quality improvement by PRS-SIM compared to Conv. SIM. PRS-SIM is capable to achieve comparable performance as GT-SIM even with the input SNR as low as ∼1 dB. Scale bar, 1 μm.

**Extended Data Fig. 3.**
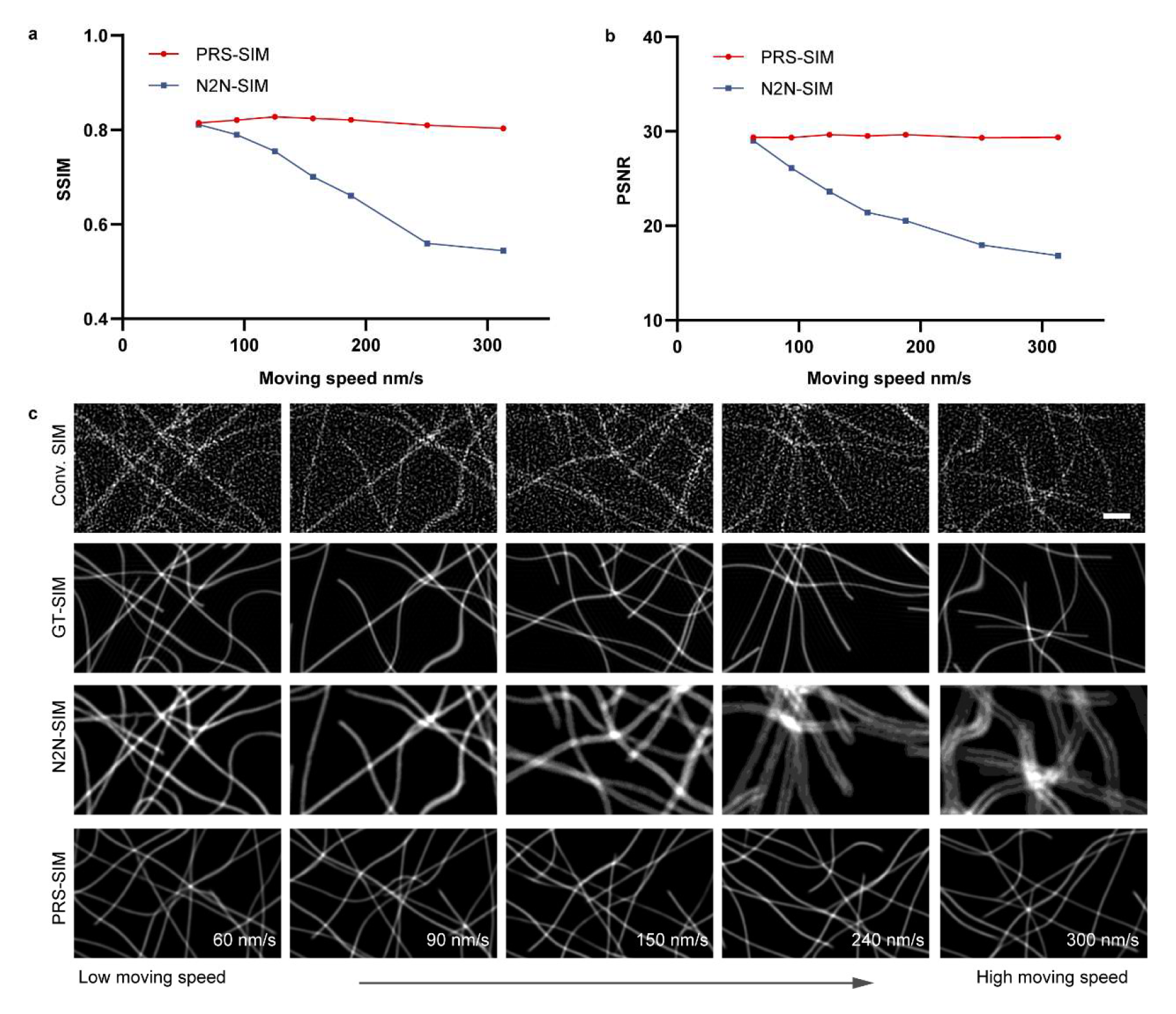
Comparison between PRS-SIM and N2N-SIM on synthetic moving microtubules. **a,b,** Quantitative comparison of PRS-SIM and N2N-SIM images in terms of PSNR and SSIM on simulated microtubules of different moving speeds. The reconstruction fidelity of N2N-SIM drops significantly as the moving speed increases, while the performance of PRS-SIM remains stable. Sample size: N=50 for each data point. **c,** Representative SR images of moving microtubules generated by Conv. SIM, GT-SIM, N2N-SIM, and PRS-SIM. The visualization results demonstrated that severe blurring artifact emerged in N2N-SIM images when the moving speed is high, while PRS-SIM is not affected since it does not rely on any temporal correlation between the adjacent frames. Scale bar, 1 μm.

**Extended Data Fig. 4.**
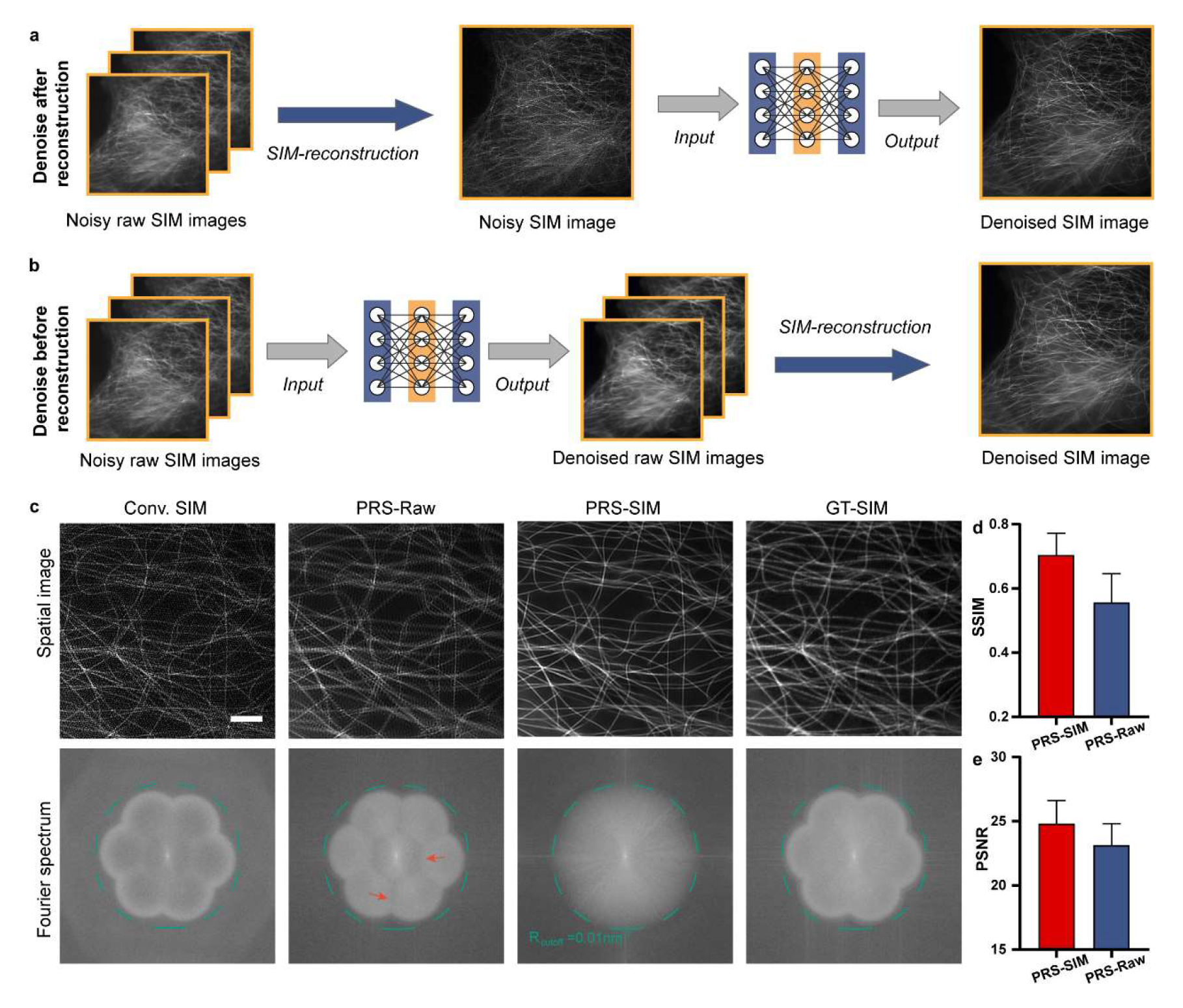
Comparison of different denoising strategies for SIM. **a,b,** The diagram of two different denoising strategies, which employ the denoising network after (**a**, denoted as PRS-SIM) and before (**b**, denoted as PRS-Raw) the conventional SIM reconstruction, respectively. **c,** Representative SIM images reconstructed with PRS-SIM and PRS-Raw and the corresponding Fourier power spectrums. The OTF cutoff frequency is annotated by dashed green circles. The PRS-Raw image contains severe ringing artifacts, which is consistent with the heterogeneous regions in its Fourier power spectrum as noted by red arrows. **d,e,** Quantitative comparison of PRS-SIM and PRS-Raw in terms of SSIM (**d**) and PSNR (**e**). Both SSIM result and PSNR results indicated that PRS-SIM achieved better performance than PRS-Raw. Sample size: N=40 for each method. Scale bar, 2 μm.

**Extended Data Fig. 5.**
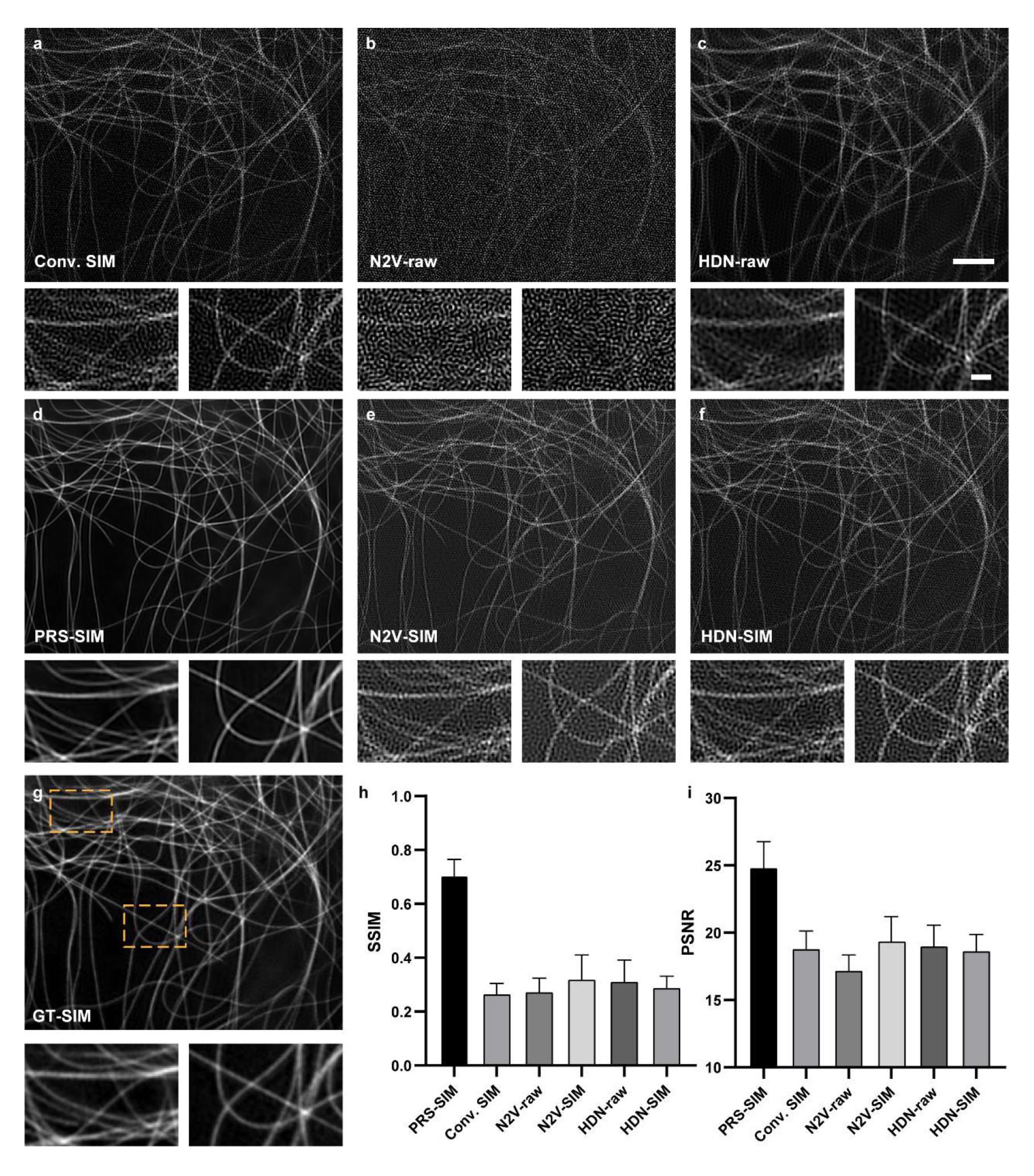
Performance comparison among different self-supervised deep-learning denoising methods. **a-f,** Representative results of the conventional SIM (**a**), PRS-SIM (**d**), noise2void (**b,e**, denoising networks employed before/after SIM reconstruction are noted as N2V-raw/N2V-SIM), hierarchical denoising network (**d,f**, denoising networks employed before/after SIM reconstruction are noted as HDN-raw/HDN-SIM). **g,** The corresponding GT-SIM image. Two zoom-in regions noted by yellow squares are shown for detailed comparison. **h,i,** Statistical comparison of the aforementioned methods by calculating the SSIM (**h**) and PSNR(**i**) referring to the GT-SIM image. The quantitative results inditated that PRS-SIM acquired the best denoising performance, which is consistent with lowest artifact and highest fidelity shown in the reconstruction images. Sample size: N=40 for each method. Scale bar, 2 μm, 0.5 μm (zoom-in regions).

**Extended Data Fig. 6.**
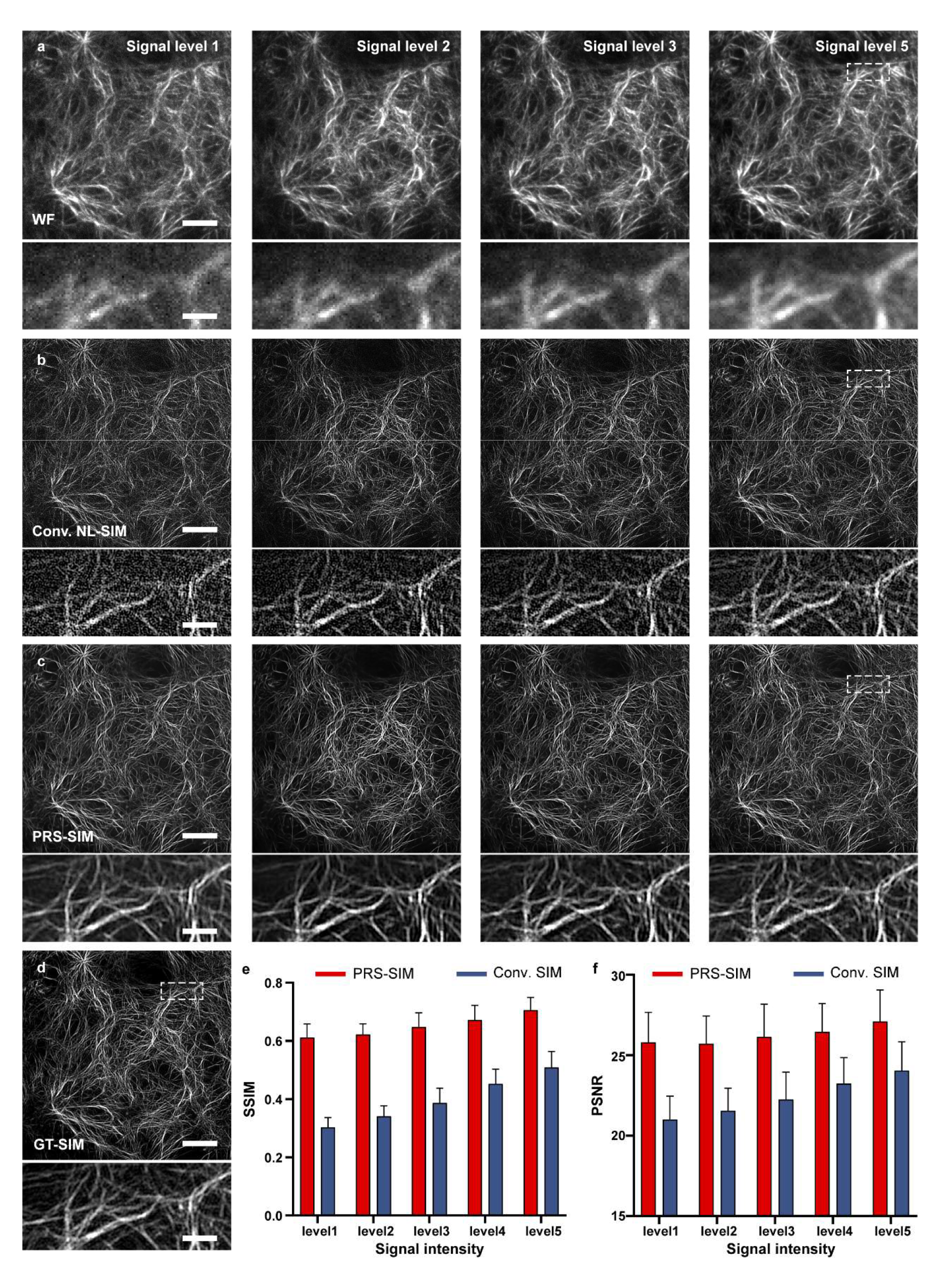
Evaluation of PRS-SIM for non-linear-SIM (NL-SIM) data denoising. **a-c,** Representative WF (**a**), conventional NL-SIM (**b**), and PRS-SIM (**c**) under different signal levels, corresponding to the same ground-truth (**d**). **e,f,** Quantitative comparison of PRS-SIM and conventional NL-SIM in terms of SSIM (**e**) and PSNR (**f**) on F-actin images. The SSIM and PSNR values referring to GT images under different signal levels are displayed. Sample size: N=20 for each signal level. Scale bar, 5 μm (regular), 1 μm (zoom-in regions).

**Extended Data Fig. 7.**
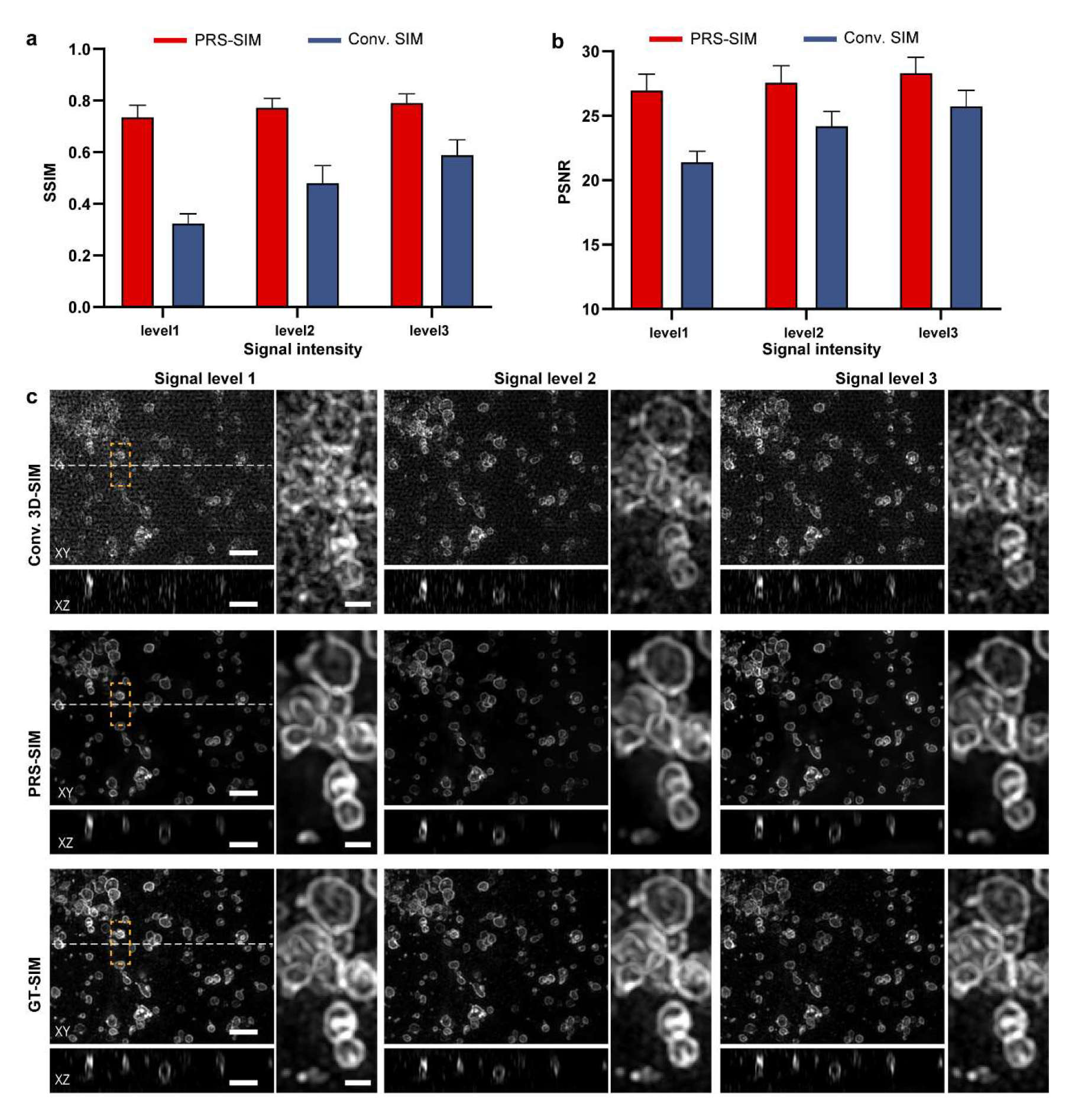
Evaluation of PRS-SIM for 3D-SIM data denoising. **a,b,** Quantitative comparison of PRS-SIM and conventional 3D-SIM in terms of SSIM (**a**) and PSNR (**b**) on Lyso images. The SSIM and PSNR values referring to GT-SIM under different signal levels are displayed. **c,** Representative images of conventional 3D-SIM, PRS-SIM, and GT SIM of different signal levels. The MIP view in XY plane and the sectioned view in XZ plane (indicated by the green line in XY view) are displayed. Compared to conventional 3D-SIM result, PRS-SIM is capable to remove most artifact in all three dimensions and achieves comparable quality and resolution to GT-SIM. Sample size: N=17 for each signal level. Scale bar, 2 μm (XY and XZ views), 0.5 μm (zoom-in regions).

**Extended Data Fig. 8.**
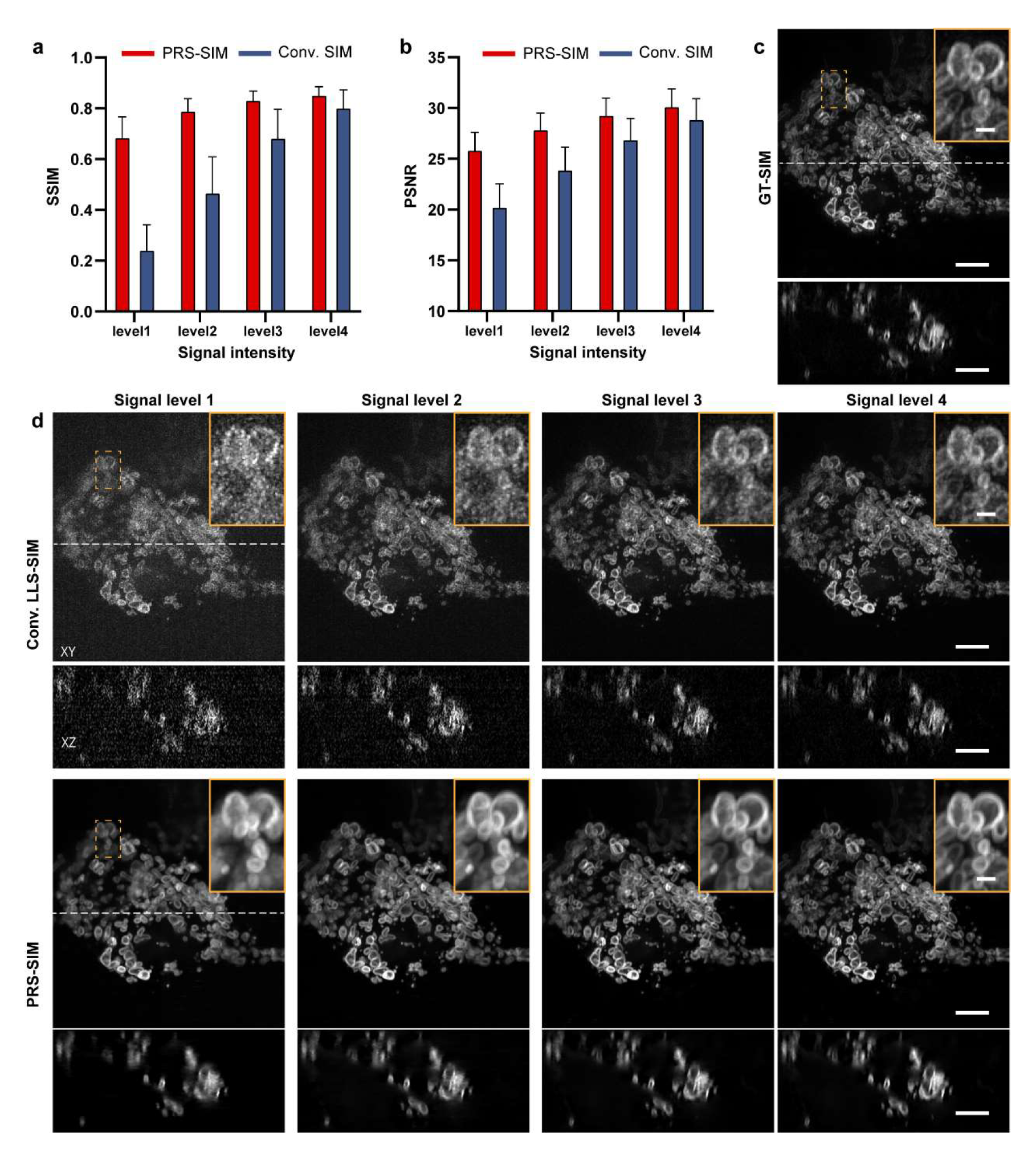
Evaluation of PRS-SIM for LLS-SIM data denoising. **a,b,** Quantitative comparison of PRS-SIM and conventional LLS-SIM in terms of SSIM (**a**) and PSNR (**b**) on ER images. The SSIM and PSNR values referring to GT images under different signal levels are displayed. **c,d,** Representative SR images of GT-SIM (**b**), conventional LLS-SIM, and PRS-SIM (**d**) under different signal levels. The MIP view in XY plane and the sectioned view in XZ plane (indicated by the green line in XY view) are displayed. Sample size: N=20 for each signal level. Scale bar, 5 μm (XY and XZ views), 1 μm (zoom-in regions).

**Extended Data Fig. 9.**
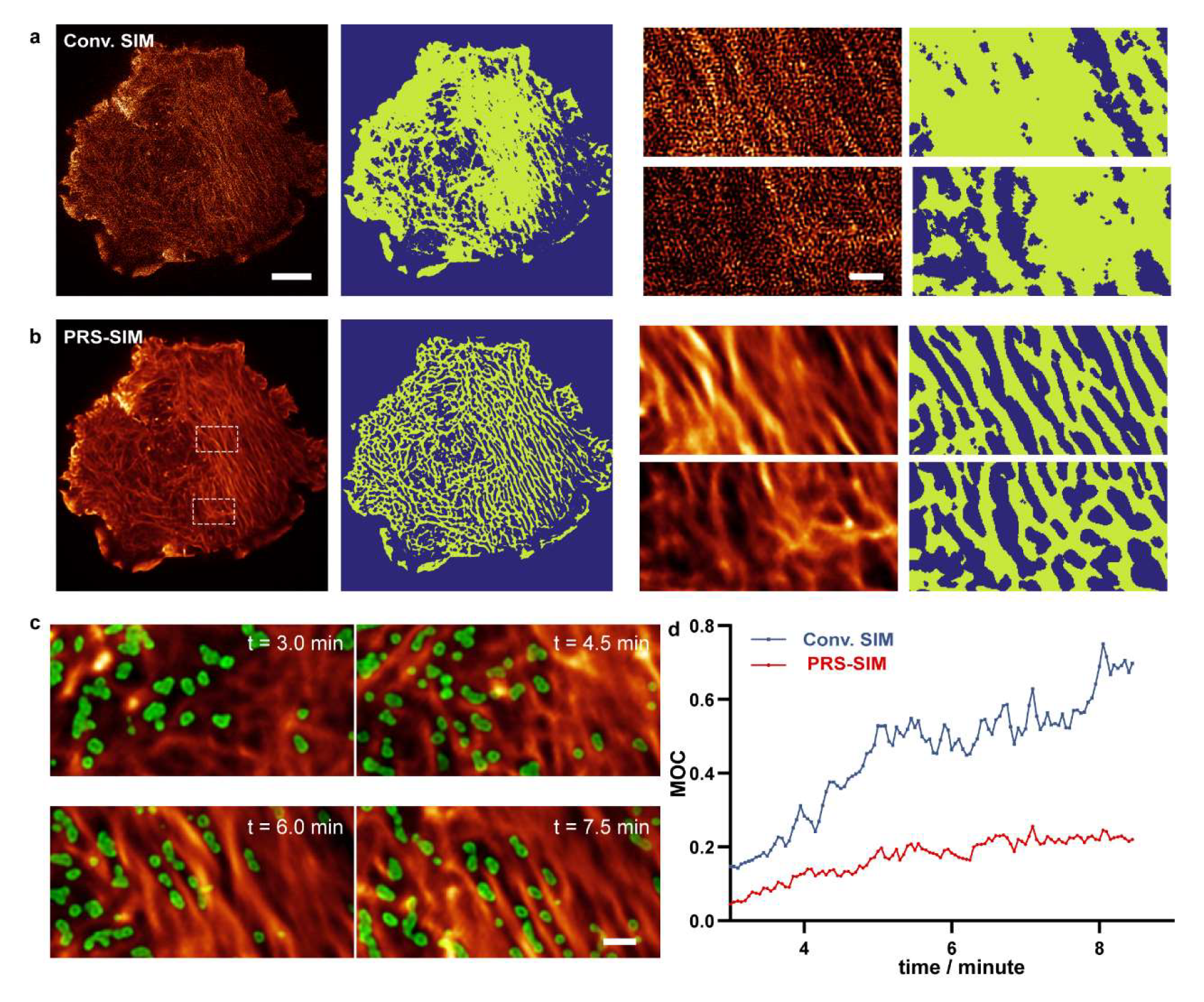
Interaction analysis of CCPs and F-actin during the growth process of a SUM159 cell via long-term PRS-SIM imaging. **a,b,** Weka segmentation results of F-actin filaments from the conventional TIRF-SIM images (**a**) and PRS-SIM images (**b**). Zoom-in views of two representative regions are displayed on the right panel. Since the experiment was performed under extremely low excitation intensity, the filaments cannot hardly be distinguished in conventional SIM images, while the quality of PRS-SIM result is adequate for successful segmentation. **c,** Representative visualization of the interaction between CCPs (green) and F-actin (red) by PRS-SIM. **d,** Mander’s overlapped coefficient of the CCPs referring to F-actin calculated from conv. SIM (blue) and PRS-SIM (red) images, respectively, during the entire cell growth process. Low MOC values indicate that most CCPs tend to locate in the interspace of actin filaments, which is consistent with the images shown in **c**. Scale bar, 5 μm (a, b), 1 μm (c, zoom-in regions in a, b).

**Extended Data Fig. 10.**
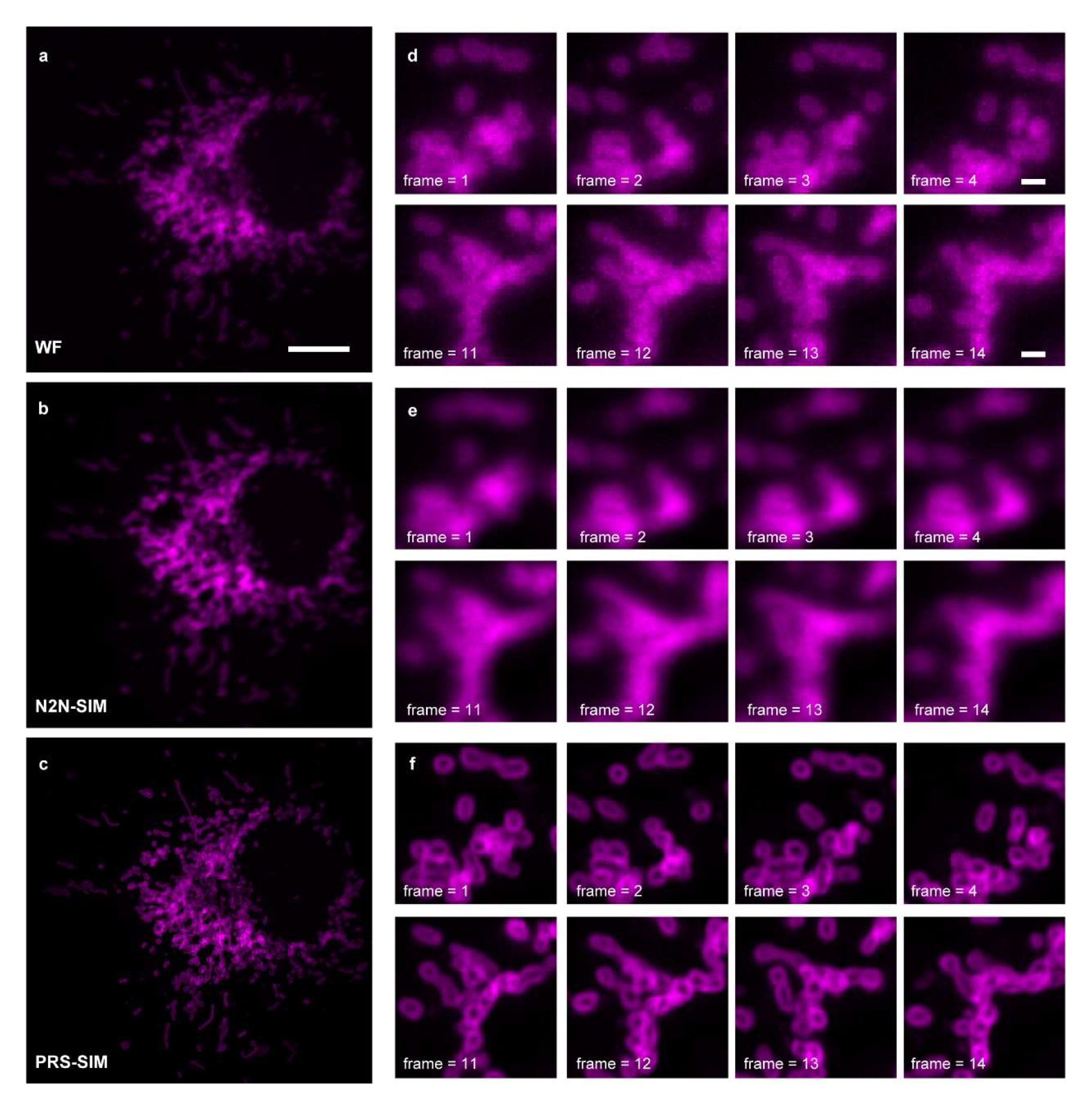
Comparison between PRS-SIM and N2N-SIM on time-lapse data of rapidly moving mitochondria. **a-c,** WF (**a**), N2N-SIM (**b**), and adaptively trained PRS-SIM (**c**) images of a COS7 cell expressing TOMM20-2xmEmerald. **d-f,** Representative time-lapse zoom-in regions of WF (**d**), N2N-SIM (**e**), and PRS-SIM (**f**) images. These results show that the temporal continuity-based N2N-SIM generates blurry artifacts because of the rapid movement of the specimen, while the proposed PRS-SIM successfully recover the fine structure of mitochondria. Scale bar, 5 μm (a-c), 0.5 μm (d-f).

