## Supplementary Information for "Self-supervised denoising for structured illumination microscopy enables long-term super-resolution live-cell imaging"

#### Abbreviations

SIM: structured illumination microscopy

PRS-SIM: pixel-realignment based self-supervised reconstruction for SIM

N2N: noise-to-noise

TIRF: total internal reflection fluorescent

WF: wide field

LLS: lattice light sheet

SR: super-resolution

HR: high-resolution

LR: low-resolution

SNR: signal-to-noise ratio

SSIM: structural similarity

PSNR: peak signal-to-noise ratio

PSF: point spread function

OTF: optical transfer function

GT: ground truth

MT: microtubule

CCP: clathrin coated pit

ER: endoplasmic reticulum

Mito: mitochondrion

Lyso: lysosome

### Supplementary Tables

#### 1. Supplementary Table 1 | Imaging condition for all live cell experiments

| | Imaging modality | Cell type | FOV | Fluorescent label | Exposure time | Cycle time | Time points | Excitation wavelength |
|---|---|---|---|---|---|---|---|---|
| Fig. 3a-b, Extended Data Fig. 8 | TIRF-SIM | SUM159 | 64um × 64um | 488: EGFP<br>560: Lifeact-Halo | 2ms<br>5ms | 3s | 170 | 488nm<br>560nm |
| Fig. 3a | TIRF-SIM | SUM159 | 64um × 64um | 488: EGFP | 2ms | 0.5s | 5000 | 488nm |
| Fig. 4a-c | 3D-SIM | COS7 | 32um × 32um × 4um | 488: 3xmEmerald-Ensconsin<br>560: Lamp1-Halo. | 5ms<br>5ms | 10s | 400 | 488nm<br>560nm |
| Fig. 4d | LLS-SIM | COS7 | 32um × 32um × 4um | 488: TOMM20-2xmEmerald<br>560: 3xmCherry-Ensconsin | 20ms | 12s | 313 | 488nm<br>560nm |
| Fig. 4e-f, Extended Data Fig. 8 | LLS-SIM | COS7 | 32um × 32um × 4um | 488: TOMM20-2xmEmerald<br>560: 3xmCherry-Ensconsin | 20ms | 12s | 151 | 488nm<br>560nm |

#### 2. Supplementary Table 2 | Training details of PRS-SIM models

| Imaging modality | Sample type | Network type | Number of raw images/volumes | Training patch size | Initial learning rate | Training batch size | Ending iteration |
|---|---|---|---|---|---|---|---|
| TIRF-SIM | CCPs | 2D | 40 | 128×128 | $1 \times 10^{-4}$ | 4 | 80000 |
| | MTs | 2D | 40 | 128×128 | $1 \times 10^{-4}$ | 4 | 80000 |
| | ER | 2D | 50 | 128×128 | $1 \times 10^{-4}$ | 4 | 80000 |
| | F-actin | 2D | 40 | 128×128 | $1 \times 10^{-4}$ | 4 | 80000 |
| 3D-SIM | MTs | 3D | 20 | 64×64×8 | $1 \times 10^{-4}$ | 4 | 100000 |
| | Lyso | 3D | 17 | 64×64×8 | $1 \times 10^{-4}$ | 4 | 100000 |
| | MTs (time lapsing) | 3D | 17 | 64×64×8 | $1 \times 10^{-4}$ | 4 | 100000 |
| | Lyso (time lapsing) | 3D | 17 | 64×64×8 | $1 \times 10^{-4}$ | 4 | 100000 |
| LLS-SIM | ER | 2D | 15 | 128×128 | $1 \times 10^{-4}$ | 4 | 100000 |
| | Mito | 2D | 20 | 128×128 | $1 \times 10^{-4}$ | 4 | 100000 |
| | MTs (time lapsing) | 2D | 16 | 128×128 | $1 \times 10^{-4}$ | 4 | 100000 |
| | Mito (time lapsing) | 2D | 16 | 128×128 | $1 \times 10^{-4}$ | 4 | 100000 |
| | MTs (N2N) | 2D-N2N | 16 | 128×128 | $1 \times 10^{-4}$ | 4 | 100000 |
| | Mito (N2N) | 2D-N2N | 16 | 128×128 | $1 \times 10^{-4}$ | 4 | 100000 |

### Supplementary Notes

#### 1. Supplementary Note 1 | Theoretical derivation of PRS-SIM

##### a. Denoising without clean data

Due to the complex environment and the photoelectric conversion instrument, detection noise is unavoidable in fluorescence imaging. Take the Gaussian white noise, which is the major type of noise in a digital image, into consideration, the detected image $\boldsymbol{y}$ can be modelled as:

$$\boldsymbol{y} = \boldsymbol{x} + \boldsymbol{n} \tag{3}$$

where $\boldsymbol{x}$ denotes the noise-free image and $\boldsymbol{n}$ denotes random noise following the normal distribution. The goal of image denoising task is to retrieve clean image $\boldsymbol{x}$ from noisy input $\boldsymbol{y}$.

Conventional image denoising algorithm mainly utilized certain analytical models to suppress the noise, such as Wiener filtering algorithm and BM3D algorithm[1]. However, the performance of these algorithms encounters the bottleneck since the sample information is strongly coupled with the random noise, especially under low SNR conditions. Benefiting from the feature extraction and representation ability of the neural network, deep-learning based algorithms have shown substantially improved performance in image denoising tasks than conventional methods[2]. For supervised denoising neural networks, the training dataset consists of a series of noisy/clean image pairs, and the objective function is defined as:

$$\min_{\varphi} \|\varphi(\boldsymbol{y}) - \boldsymbol{x}\|_2^2, \tag{4}$$

where $\varphi(\cdot)$ denotes the neural network.

However, for most biological samples, the acquisition of clean GT data is difficult and costly. To this end, in recent years several weakly supervised or self-supervised denoising methods, which do not require clean GT data, have aroused increasing interest. Among these methods, Noise2noise[3] (N2N) is one of the most representative techniques, which employs two independently captured noisy images of the same sample as the input and target during network training. The underlying mechanism of N2N is that the mathematical expectation of its optimization function is equivalent to that of supervised learning described in Eq. (4). Let $\boldsymbol{y_1} = \boldsymbol{x} + \boldsymbol{n_1}$ and $\boldsymbol{y_2} = \boldsymbol{x} + \boldsymbol{n_2}$ denotes the two independently captured images, where $\boldsymbol{n_1}$ and $\boldsymbol{n_2}$ following normal distribution $N(0, \sigma^2)$. The mathematical expectation of objective function of N2N is formulated as:

$$
\begin{aligned}
E\{\|\varphi(\boldsymbol{y_1})-\boldsymbol{y_2}\|_2^2\} &= E\{\|\varphi(\boldsymbol{y_1})-\boldsymbol{x}+\boldsymbol{n_2}\|_2^2\}\\
&= E\{\|\varphi(\boldsymbol{y_1})-\boldsymbol{x}\|_2^2\}-2E\{(\varphi(\boldsymbol{y_1})-\boldsymbol{x})\boldsymbol{n_2}\}+E\{\boldsymbol{n_2^2}\}\\
&= E\{\|\varphi(\boldsymbol{y_1})-\boldsymbol{x}\|_2^2\}-2E\{\varphi(\boldsymbol{y_1})-\boldsymbol{x}\}E\{\boldsymbol{n_2}\}+E\{\boldsymbol{n_2^2}\}\\
&= E\{\|\varphi(\boldsymbol{y_1})-\boldsymbol{x}\|_2^2\}+E\{\boldsymbol{n_2^2}\}\\
&= E\{\|\varphi(\boldsymbol{y_1})-\boldsymbol{x}\|_2^2\}+\sigma^2 \qquad (5)
\end{aligned}
$$

The derivation of Eq. (5) utilized the truth that $E\{\boldsymbol{n_1}\}=E\{\boldsymbol{n_2}\}=0$ (zero-mean) and $2E\{(\varphi(\boldsymbol{y_1})-\boldsymbol{x})\boldsymbol{n_2}\}=2E\{\varphi(\boldsymbol{y_1})-x\}E\{\boldsymbol{n_2}\}$ (statistical independence). Eq. (5) shows that the expectation of the loss function defined on $\boldsymbol{y_1}$ and $\boldsymbol{y_2}$ is equivalent to the supervised one defined on $\boldsymbol{y}$ and $\boldsymbol{x}$ except for a constant. Because of its ingenious theory and superior denoising performance, N2N has become a millstone algorithm and enlightened several subsequent denoising schemes[4].

#### b. Self-supervised denoising with similar scenario

Although N2N has shown great denoising performance on either natural images or microscopic images, the requirement of duplicated captures of the same specimen strictly limits its application. To further develop denoising techniques applicable for single-captured SIM images, we investigated data augment strategy termed neighbor2neighbor[5], which exploiting the similarity between adjacent pixels to create the training dataset. Specifically, let $\boldsymbol{y_A}$ and $\boldsymbol{y_B}$ denote two sub-images extracted from the same noisy image $\boldsymbol{y}$ corresponding to the ground-truth $\boldsymbol{x}$, it is easy to prove that the noise in $\boldsymbol{y_A}$ and $\boldsymbol{y_B}$ follows the independent zero-mean Gaussian distribution. So their conditional mathematical expectation is represented as:

$$E_{\boldsymbol{y_A}|\boldsymbol{x}}(\boldsymbol{y_A})=\boldsymbol{x} \qquad (6)$$

$$E_{\boldsymbol{y_A}|\boldsymbol{x}}(\boldsymbol{y_A})=\boldsymbol{x}+\boldsymbol{\varepsilon} \qquad (7)$$

where $\boldsymbol{\varepsilon}$ denotes the difference between the underlying ground-truth of $\boldsymbol{y_A}$ and $\boldsymbol{y_B}$.

$$E_{\boldsymbol{y_A}|\boldsymbol{x}}(\|\varphi(\boldsymbol{y_A})-\boldsymbol{x}\|_2^2)=E_{\boldsymbol{y_A},\boldsymbol{y_B}|\boldsymbol{x}}(\|\varphi(\boldsymbol{y_A})-\boldsymbol{y_B}+\boldsymbol{y_B}-\boldsymbol{x}\|_2^2)$$

$$=E_{\boldsymbol{y_A}|\boldsymbol{x}}(\|\varphi(\boldsymbol{y_A})-\boldsymbol{y_B}\|_2^2)+E_{\boldsymbol{y_B}|\boldsymbol{x}}(\|\boldsymbol{y_B}-\boldsymbol{x}\|_2^2)+2E_{\boldsymbol{y_A},\boldsymbol{y_B}|\boldsymbol{x}}(\varphi(\boldsymbol{y_A})-\boldsymbol{y_B})\cdot 2E(\boldsymbol{y_B}-\boldsymbol{x})$$

$$=E_{\boldsymbol{y_A}|\boldsymbol{x}}(\|\varphi(\boldsymbol{y_A})-\boldsymbol{y_B}\|_2^2)+\sigma_n^2+2\boldsymbol{\varepsilon}\cdot E_{\boldsymbol{y_A},\boldsymbol{y_B}|\boldsymbol{x}}(\varphi(\boldsymbol{y_A})-\boldsymbol{y_B}), \qquad (8)$$

where $\sigma_n^2$ denotes the covariance of the noisy image $\boldsymbol{y_B}$.

Although $\boldsymbol{\varepsilon}$ is a non-zero variable, since the sub-images $\boldsymbol{y_A}$ and $\boldsymbol{y_B}$ is extracted by the adjacent pixels, the exact value of $\boldsymbol{\varepsilon}$ should be comparable small, so that the item $2\boldsymbol{\varepsilon} \cdot E_{\boldsymbol{y_A},\boldsymbol{y_B}|\boldsymbol{x}}(\varphi(\boldsymbol{y_A}) - \boldsymbol{y_B})$ is close to zero. As an acceptable approximation, $E_{\boldsymbol{y_A}|\boldsymbol{x}}(\|\varphi(\boldsymbol{y_A}) - \boldsymbol{x}\|_2^2)$ can be represented as:

$$E_{\boldsymbol{y_A}|\boldsymbol{x}}(\|\varphi(\boldsymbol{y_A}) - \boldsymbol{x}\|_2^2) \approx E_{\boldsymbol{y_A},\boldsymbol{y_B}|\boldsymbol{x}}(\|\varphi(\boldsymbol{y_A}) - \boldsymbol{y_B}\|_2^2) + \sigma_n^2 \tag{9}$$

Since $E_{\boldsymbol{y_A},\boldsymbol{x}}(\cdot) = E_{\boldsymbol{y_A}|\boldsymbol{x}}(\cdot)E_{\boldsymbol{x}}(\cdot)$, we further have:

$$E_{\boldsymbol{y_A},\boldsymbol{x}}(\|\varphi(\boldsymbol{y_A}) - \boldsymbol{x}\|_2^2) \approx E_{\boldsymbol{y_A},\boldsymbol{y_B},\boldsymbol{x}}(\|\varphi(\boldsymbol{y_A}) - \boldsymbol{y_B}\|_2^2) + \sigma_n^2 \tag{10}$$

In Eq. (10), the item $E_{\boldsymbol{y_A},\boldsymbol{y_B},\boldsymbol{x}}(\|\varphi(\boldsymbol{y_A}) - \boldsymbol{y_B}\|_2^2)$ is the mathematical expectation of the loss function to train the network utilizing $\boldsymbol{y_A}$ and $\boldsymbol{y_B}$ as the input and the target, respective, the item $E_{\boldsymbol{y_A},\boldsymbol{x}}(\|\varphi(\boldsymbol{y_A}) - \boldsymbol{x}\|_2^2)$ is the loss function of the supervised training, and their difference between these two loss functions is only a constant. Therefore, as long as the underlying ground-truth between $\boldsymbol{y_A}$ and $\boldsymbol{y_B}$ is similar, to train the network $\boldsymbol{y_A}$ and $\boldsymbol{y_B}$ is approximated to supervised training with clean data. Since $\boldsymbol{y_A}$ and $\boldsymbol{y_B}$ is generated only from a single noisy image $\boldsymbol{y}$, these scheme is an effective self-training scheme without the requirement of clean data or repeatedly acquisition of the same scene as used in N2N[3].

**c. Pixel-realignment strategy for SIM denoising**

After introducing the self-supervised denoising strategy with similar scenario, we make a further step in this section to incorporate it with SIM reconstruction and denoising. Before proceeding, we take a brief re-visit to the convention SIM algorithm with the mixed Poisson and Gaussian noise assumption. The super-resolution (SR) SIM image is reconstructed from a series of raw SIM images with different illumination patterns. Ignoring the non-linear effect of the fluorescent probe, the emission intensity is proportional to excitation, so that the detected image $D(\boldsymbol{r})$ can be represented as:

$$D(\boldsymbol{r}) = [S(\boldsymbol{r}) \cdot I(\boldsymbol{r})] \otimes PSF(\boldsymbol{r}) + N(\boldsymbol{r}), \tag{11}$$

where $S(\boldsymbol{r})$ represents the ground-truth of the sample, $I(\boldsymbol{r})$ denotes the spatial illumination pattern, $PSF(\boldsymbol{r})$ is the point spread function (PSF) of the imaging system and $N(\boldsymbol{r})$ denotes the detection noise. The typical sinusoidal illumination pattern used in SIM can be mathematically represented as the sum of several sinusoidal components:

$$I(\boldsymbol{r}) = \sum_{m} \exp(i \cdot 2\pi \cdot m\boldsymbol{p} \cdot \boldsymbol{r} + m\phi) \tag{12}$$

where $\boldsymbol{p}$ denotes the modulation vector, $\phi$ denotes the modulation phase, and $m$ is the order of the components, respectively.

By applying Fourier transformation, the detected image in Fourier domain is represented as:

$$\begin{aligned}
\widetilde{D}_{p,\phi}(\boldsymbol{k_r}) &= \left[\tilde{S}(\boldsymbol{k_r}) \otimes \tilde{I}(\boldsymbol{k_r})\right] \cdot OTF(\boldsymbol{k_r}) + \widetilde{N}(\boldsymbol{k_r}) \\
&= [\sum_{m} \exp(-i \cdot m\phi)\, \tilde{S}(\boldsymbol{k_r} - m\boldsymbol{p})] \cdot OTF(\boldsymbol{k_r}) + \widetilde{N}(\boldsymbol{k_r}) \\
&= \sum_{m} \exp(-i \cdot m\phi)\, \widetilde{D}_{p,m}(\boldsymbol{k_r}) + \widetilde{N}(\boldsymbol{k_r}),
\end{aligned} \tag{13}$$

where the topmark $\sim$ denotes the variables in Fourier domain, $OTF(\boldsymbol{k_r})$ is the optical transfer function of the system, and $\widetilde{D}_{p,m}(\boldsymbol{k_r}) = \tilde{S}(\boldsymbol{k_r} - m\boldsymbol{p}) \cdot OTF(\boldsymbol{k_r})$ represents the Fourier spectrum information of order $m$.

From Eq. (13), $\widetilde{D}_{p,\phi}(\boldsymbol{k_r})$ is actually the linear combination of $\widetilde{D}_{p,m}(\boldsymbol{k_r})$ added with noise. For each modulation vector $\boldsymbol{p}$, a certain number of raw SIM images, i.e., 3 for 2D-SIM and 5 for 3D-SIM, with different phase $\phi_m$ were collected. Then $\widetilde{D}_{p,m}(\boldsymbol{k_r})$ can be retrieved by the separation matrix as:

$$\begin{bmatrix} \widetilde{D}_{p,\phi_1}(\boldsymbol{k_r}) \\ \widetilde{D}_{p,\phi_2}(\boldsymbol{k_r}) \\ \cdots \\ \widetilde{D}_{p,\phi_m}(\boldsymbol{k_r}) \end{bmatrix} = \begin{bmatrix} \exp(-i \cdot m\phi_1) & \cdots & 1 & \cdots & \exp(i \cdot m\phi_1) \\ \exp(-i \cdot m\phi_2) & \cdots & 1 & \cdots & \exp(i \cdot m\phi_2) \\ \cdots & \cdots & \cdots & \cdots & \cdots \\ \exp(-i \cdot m\phi_m) & \cdots & 1 & \cdots & \exp(i \cdot m\phi_m) \end{bmatrix} \begin{bmatrix} \widetilde{D}_{p,-m}(\boldsymbol{k_r}) \\ \widetilde{D}_{p,-m+1}(\boldsymbol{k_r}) \\ \cdots \\ \widetilde{D}_{p,m}(\boldsymbol{k_r}) \end{bmatrix}. \tag{14}$$

For each separated spectrum component $\widetilde{D}_{p,m}(\boldsymbol{k_r})$, we shifted it to the corresponding region in the Fourier domain, applied an generalized Wiener filter to stitch them together and employed an apodization filter to mimic the real experimental OTF, then the final reconstructed SR-SIM image was formulated as:

$$\tilde{\tilde{S}}(\boldsymbol{k_r}) = \frac{\sum_{p,m} \widetilde{D}_{p,m}(\boldsymbol{k_r} - m \cdot \boldsymbol{p}) \cdot OTF(\boldsymbol{k_r} - m\boldsymbol{p})}{\|OTF(\boldsymbol{k_r} - m \cdot \boldsymbol{p})\|^2 + \omega^2} A(\boldsymbol{k_r}), \tag{15}$$

where $\omega$ denotes the filtering parameter and $A(\boldsymbol{k_r})$ is an apodization function to make the expanded OTF close to the real OTF.

Based on previous discussion, all the data processing steps in conventional SIM algorithm, including the (inverse) Fourier transformation, Wiener filtering, and matrix multiplication, are linear operations, so that the final constructed SR images can be represented as a linear combination of all raw SIM images. We used $\boldsymbol{y_1}, \boldsymbol{y_2}, \cdots \boldsymbol{y_N}$ denoting the raw SIM image sequence, where $N$ denotes the image number of each SIM image group, i.e., $3 \times 3$ for 2D/TIRF-SIM, $3 \times 5$ for 3D-SIM and $1 \times 3$ for LLS-SIM. The final reconstructed SR $\boldsymbol{Y}$ image can be represented as:

$$\boldsymbol{Y} = \sum_{i=1}^{N} c_i \boldsymbol{y_i}, \quad (16)$$

where $c_i$ denotes the weights for each raw image.

From the raw SIM image group, the pixel realignment strategy is implemented as:

a) Divide each each raw image $\boldsymbol{y_i}$, into 4 sub-images $\boldsymbol{y_{i,A}}$, $\boldsymbol{y_{i,B}}$, $\boldsymbol{y_{i,C}}$, $\boldsymbol{y_{i,D}}$ by applying a $2 \times 2$ down-sampler and form four sub-image groups.
b) Up sample each sub-image group at a factor of 2 into the original size with the nearest interpolation.
c) Apply a sub-pixel translation to each image, based on the position of the valid pixel in each $2 \times 2$ cell to guarantee them in the same spatial position.
d) Reconstructed the sub-image groups with conventional SIM algorithm and generate 4 noisy SIM SR images $\boldsymbol{Y_A}$, $\boldsymbol{Y_B}$, $\boldsymbol{Y_C}$ and $\boldsymbol{Y_D}$.
e) Randomly select 2 out of 4 noisy SIM images as the input and target image, respectively.

Based on previous discussion, the final selected SR images (e.g. $\boldsymbol{Y_A}$ and $\boldsymbol{Y_B}$) are represented as:

$$\boldsymbol{Y_A} = \sum_{i=1}^{N} c_i TU(\boldsymbol{y_{i,A}}), \quad (17)$$

$$\boldsymbol{Y_B} = \sum_{i=1}^{N} c_i TU(\boldsymbol{y_{i,B}}), \quad (18)$$

where $TU(\cdot)$ represents the integrated operator for up-sampling and translation. From Eq. (17) and Eq. (18), since all the operations are linear without offset and the noise in each raw image $\boldsymbol{y_{i,A}}$ and $\boldsymbol{y_{i,B}}$ is independent, the noise in $\boldsymbol{Y_A}$ and $\boldsymbol{Y_B}$ follows independent zero-mean Gaussian distribution. Moreover, since each raw image $\boldsymbol{y_{i,A}}$ and $\boldsymbol{y_{i,B}}$ shares the similar ground-truth, the underlying gap between in $\boldsymbol{Y_A}$ and $\boldsymbol{Y_B}$ is also in an acceptable range. So to train the network with $\boldsymbol{Y_A}$ and $\boldsymbol{Y_B}$ satisfies the simularity criteria, the effectiveness of PRS-SIM is proved.

#### 2. Supplementary Note 2 | Simulation of PRS-SIM

To quantitatively evaluate the performance of PRS-SIM, we built a simulation model to synthetic raw SIM data. To make the simulated model closer to the experimental conditions, we used identical imaging parameters with our TIRF-SIM system: detection NA = 1.3, excitation NA = 1.17, excitation wavelength = 488 nm, and emission wavelength = 525 nm. The orientations of the three illumination patterns are 1.58, 2.62 and 0.53 rad and the phases used for each orientation are 0, $2/3\pi$, and $-2/3\pi$. The pixel size and shape of the raw SIM image is 62.6 nm and 256×256, respectively. We generated the structure illumination pattern and system PSF based on above settings via MATLAB and simulated the raw SIM images following the optical imaging model:

$$I_{LR}(\boldsymbol{r}) = |[Sample(\boldsymbol{r}) \cdot Pattern(\boldsymbol{r})] \otimes PSF(\boldsymbol{r})|_{\downarrow 2}, \tag{19}$$

where $Sample(\boldsymbol{r})$ denotes the grouth truth and $\downarrow 2$ represents the down-sampling operator. To mimic the detection noise, additional Gaussian white noise with $\sigma = 4.5$ and signal-related Poission noise with $\gamma$=2 were further added to the simulated wide-filed images as follows:

$$I_{detect}(\boldsymbol{r}) = \gamma \cdot P\left(\frac{I_{LR}(\boldsymbol{r})}{\gamma}\right) + N(0, \sigma^2). \tag{20}$$

The SNR of each synthetic image is defined by the ratio of the maximum signal intensity to the standard deviation of the noise. We used tubulin filaments as a representative biological structure in the simulation model. Each GT image consists 30~40 filaments with random average intensity and spatial positions in a field of view of $16\mu\text{m} \times 16\mu\text{m}$. The derived GT image was then distorted by a random elastic deformation field.[6]

We then synthetized time-lapsed image sequence of moving tubulins to mimic the live-cell imaging of organelles (Extended Data Fig. 3). The initial frame is created with the same procedure as discussed before. Then for the each following frame, a displacement of a randomly orientation relative to previous frame was assigned for each individual filament. The length of displacement $D$ is determined by the moving speed as:

$$D = \frac{1+u}{2} v_{max}, \tag{21}$$

where $v_{max}$ denotes designated the maximum moving and $u$ denotes a random variable follows the uniform distribution of $[0,1]$. The spatial shift of the filament is implemented by the *imwarp* function in MATLAB.

#### 3. Supplementary Note 3 | Fiji plugin of PRS-SIM

Deep-learning based image denoising and restoration algorithms have shown great advances in improving the quality of microscopy images. However, most of them require specific software environments such as Tensorflow or Pytorch, which can be only operated with a Python script. To make the proposed PRS-SIM more convenient for users, especially for biological researchers, we developed an easy-to-used Fiji plugin, which embeds both network training and on-click denoising functionalities. Our plugin is developed based on the open-source JAVA deep-learning framework CSBDeep (http://csbdeep.bioimagecomputing.com/) and Bioimage model zoo[7]. In this section, we will briefly introduce the workflow of PRS-SIM Fiji plugin, and more detailed instructions can be found in Supplementary software or our website (https://intellm-thuibcs.github.io/PRS-SIM).

##### a. Training with plugin

The PRS-SIM Fiji plugin is developed based on Tensorflow v.1.15, CUDA 10.1 and cuDNN 7.5. After the environment configuration and installation of the plugin, the training process mainly takes following steps (Supplementary Fig. 3):

(1) Start PRS-SIM plugin by clicking: *Fiji menu -> Plugins -> PRS-SIM -> train*.

(2) Designate the training parameters, including the total epoch number, the iteration number for each epoch, the batch size, and the initial learning rate.

(3) Click the *OK* button and the training process will begin. The training loss and validation loss will be real-time displayed in an information window.

(4) During the training process, the intermediated validation images can be visualized by clicking *preview* button. The early stop can be executed by clicking the *Finish* button.

(5) When the training is finished, click *Export Mode* button and the detailed training information window will be displayed. The trained model is saved to a designated folder by clicking *File actions -> Save to...*

##### b. Denoising with Fiji plugin

The main workflow to performance denoising with our plugin is schematized in Supplementary Fig. 4. For a noisy SIM raw image group, we firstly need to applied conventional SIM algorithm to acquire the noisy SR SIM image, which can be implemented by either our Matlab codes or the open-source software fairSIM[8], HifiSIM[9], or OpenSIM[10]. Then the inference process takes the following five steps:

(1) Open the reconstructed SIM SR image in a new window with Fiji.

(2) Start PRS-SIM plugin by clicking: *Fiji menu -> Plugins -> PRS-SIM -> predict*.

(3) Designate the path of the pre-trained model.

(4) Specify the tiling parameters including the number of tiles and the overlap between the adjacent tiles.

(5) Click the *OK* button, then the inference process will begin. The detailed information and the final denoised image will be displayed in a new Fiji window.

The inference of PRS-SIM is implemented in a tile-by-tile mode, as the input image is firstly divided into several tiles, then processed one by one, and finally stitched together to form the denoised image. To eliminate the stitching artifact, adequate overlaps between adjacent tiles are needed. Therefore, the tile size and the overlap ratio should be set properly to balance the computation resource and the data processing speed.

### Supplementary Figures

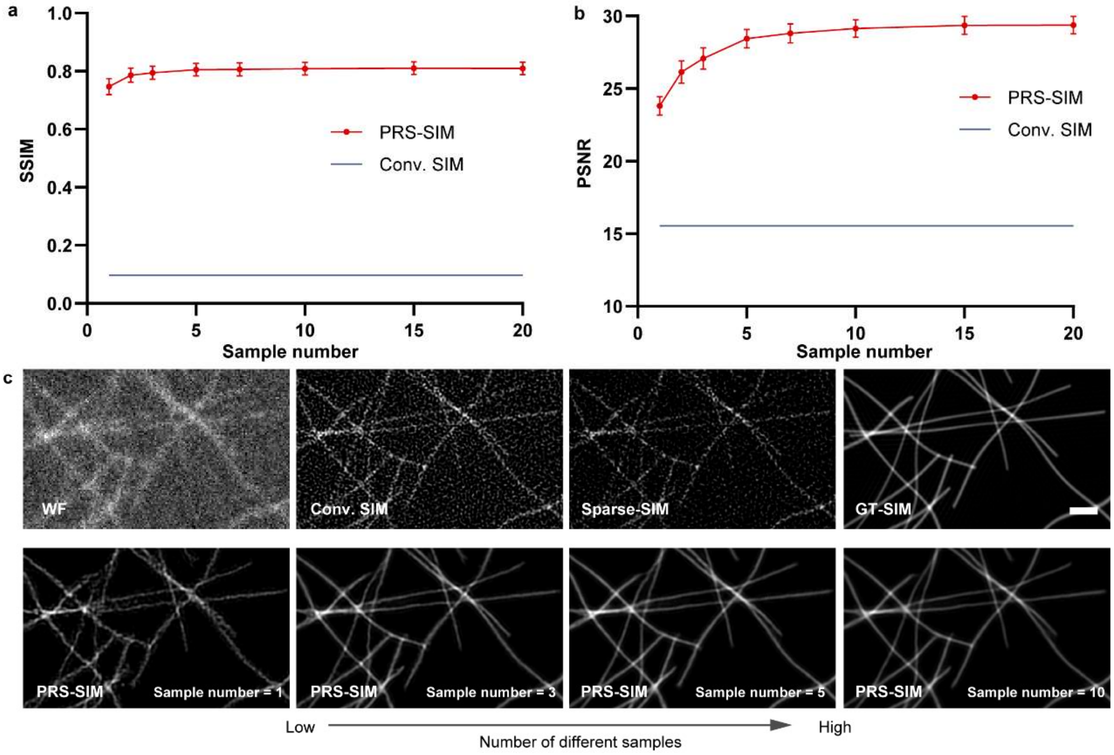

**Supplementary Fig. 1 | Evaluation of PRS-SIM models trained on different training dataset scales. a,b,** Quantitative evaluation of PRS-SIM models trained on dataset of different sample numbers. SSIM (**a**) and PSNR (**b**) are used to evaluate the performance of PRS-SIM. Sample size: N = 50 for each data point. **c,** Representative PRS-SIM denoised images trained on dataset of different sample numbers. The WF, Conv. SIM, Sparse-SIM and GT-SIM images are provided for compared. Each PRS-SIM model is trained of the specific number (as indicated) of randomly generated noisy images. Scale bar, 1μm.

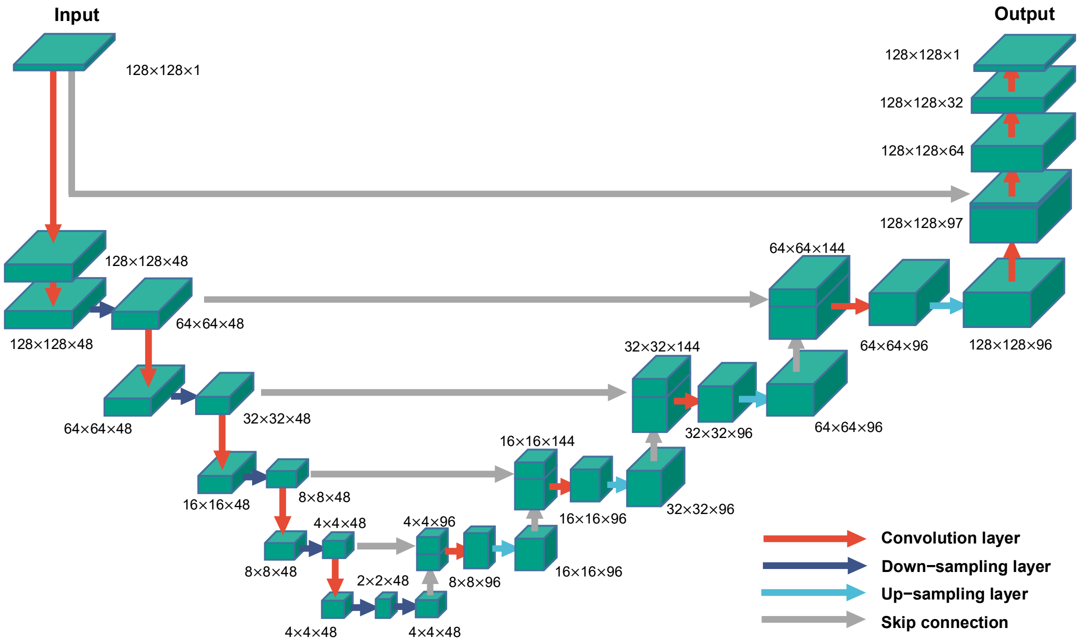

**Supplementary Fig. 2 | Network architecture.** PRS-SIM employed a U-shape convolutional neural network (U-net[11]) for image denoising. The Input image is firstly down sampled by the encoder module for feature extraction and then up sampled to the original scale by the decoder module. Skip connection are embedded to improve the training performance. During both training and inference stage, the PRS-SIM model takes noisy SIM images as inputs and output noise-free SR-SIM images.

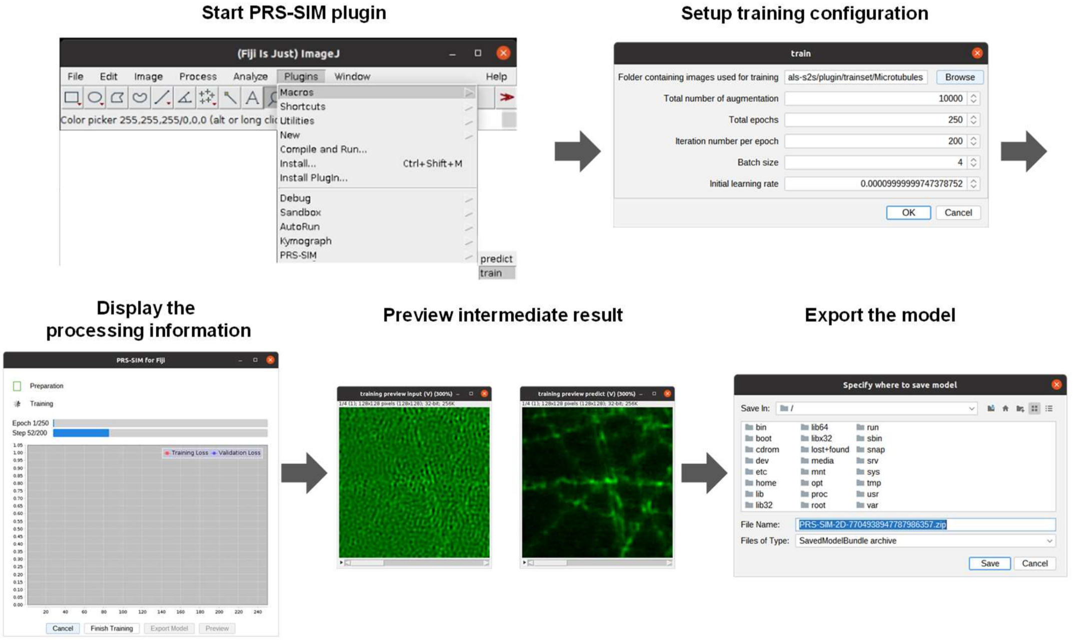

**Supplementary Fig. 3 | Network training with PRS-SIM plugin.** To make PRS-SIM convenient for users, we built an easy-to-use Fiji plugin. To train the denoising network with the PRS-SIM Fiji plugin takes the following steps: (1) Start the plugin in Fiji menu with *train* option; (2) Designate the training parameters, including the directory of the training set, initial learning rate, epoch number, etc; (3) Display the detailed training information. (4) Preview the intermediate denoising result. (5) Export the model after the training is finished.

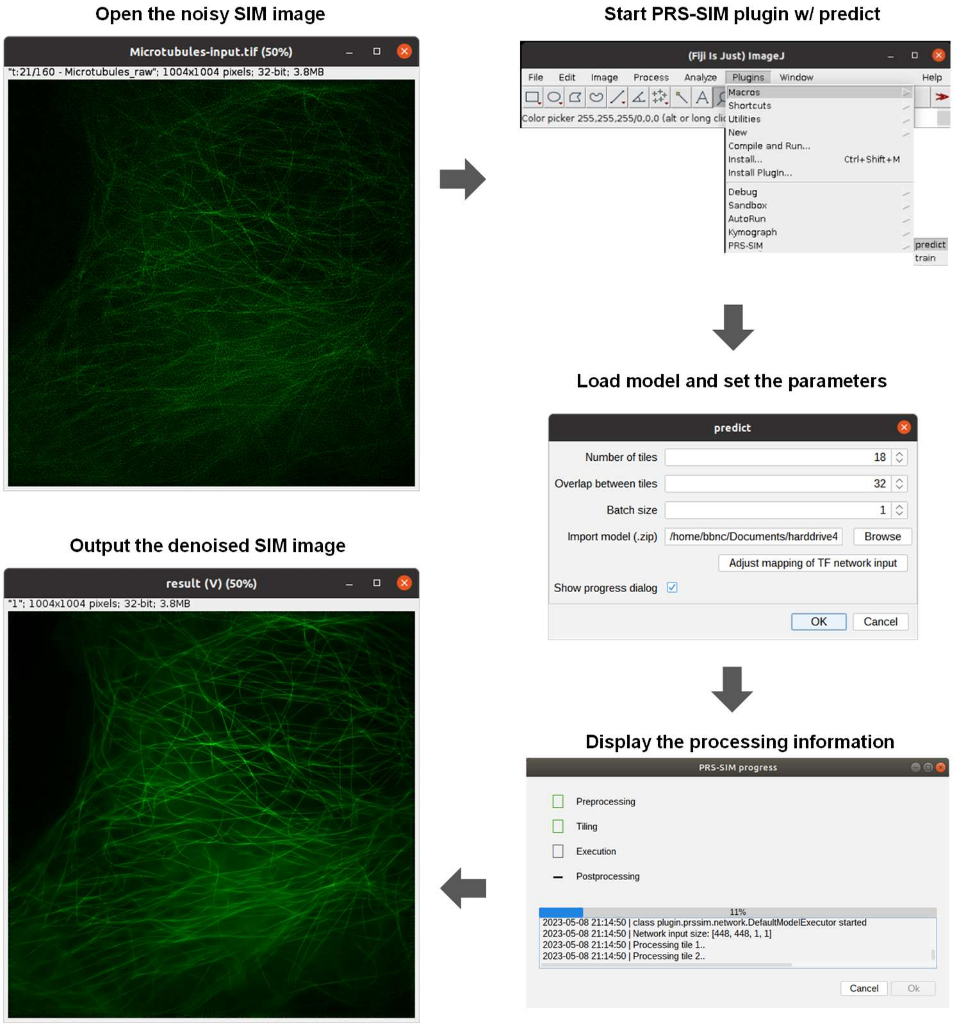

**Supplementary Fig. 4 | Denoising with PRS-SIM plugin.** The image denoising can be easily implemented with our Fiji plugin by several clicks, including: (1) Open the noisy SIM SR image to be processed; (2) Start the plugin in Fiji menu with *predict* option; (3) Load the pre-trained model file and designate the inference parameters. Then the final denoised image will be displayed in a Fiji image window accompanied with the detailed logs shown in the information window.
